# Rub1/NEDD8, a ubiquitin-like modifier, is also a ubiquitin modifier

**DOI:** 10.1101/2020.06.18.159145

**Authors:** Sylvia Zerath Gurevich, Abhishek Sinha, Joseph Longworth, Rajesh K. Singh, Betsegaw E. Lemma, Anita Thakur, Oliver Popp, Daniel Kornitzer, Noa Reis, Martin Scheffner, Gunnar Dittmar, Elah Pick, David Fushman, Michael H. Glickman

**Affiliations:** Faculty of Biology, Technion Israel Institute of Technology, Haifa 32000 Israel; Department of Chemistry and Biochemistry, University of Maryland, College Park, MD 20742, USA; Department of Biology and Environment, University of Haifa at Oranim, Tivon 36006, Israel; Max-Delbruck Center, Berlin Germany; LIH Luxemburg; Faculty of Medicine, Technion-IIT, Haifa 32000 Israel; Department of Biology - Universität Konstanz, Germany

**Keywords:** Ubiquitin, polyubiquitin, Rub1, NEDD8, Ubiquitin-like (UBL), CULLIN, Cullin-related ligases (CRL), protein post-translational modifications (PTMs)

## Abstract

Of all ubiquitin-like small protein modifiers, Rub1/NEDD8 is the closest kin of ubiquitin in sequence and in structure. Despite their profound similarities, prevalence of ubiquitin and of Rub1 is starkly different: targets of ubiquitin modification reach into the thousands, whereas unique targets of Rub1/NEDD8 appear limited to one family of proteins, Cullins. This distinction is likely due to dedicated E1 activating enzymes that select either one or the other and relay the modifier until it is covalently attached to a target. To convert typical neddylation targets for modification by ubiquitin, and vice versa, we designed reciprocal substitutions at position 72 of Rub1 and of ubiquitin to render them substrates for activation by their non-cognate E1 activating enzymes. We found that this single amino acid is sufficient to distinguish between Ub and Rub1 in living cells, and determine their targets. Thus, modification of Cullins by Ub^R72T^ could compensate for loss of Rub1, even as it maintained its ability to polymerize and direct conjugates for degradation. Conversely, Rub1^T72R^ activated by ubiquitin-activating enzyme entered into the ubiquitination cascade, however was not efficiently polymerized, essentially capping polyubiquitin chains. Upon shortage of free ubiquitin under stress, even native Rub1 spilled-over into the ubiquitinome suppressing polyubiquitination. By contrast, the need to maintain monomeric modifications on unique targets is a likely explanation for why the Rub1-activating enzyme strictly discriminates against ubiquitin. Swapping Rub1 and ubiquitin signals uncovered a reason for maintaining two separate pathways across eukaryotic kingdom.

## INTRODUCTION

The ubiquitin-like (UBL) family of small protein modifiers is a class of evolutionary conserved proteins that are reversibly conjugated to other proteins to regulate a variety of fundamental cellular processes (1-3). Ubiquitin (Ub) is the most prevalent UBL, and ubiquitination is one of the most active metabolic pathways with hundreds of enzymes involved in ubiquitination, recognition, or deubiquitination of thousands of target proteins (4-6). By contrast, unique targets of Rub1 (Related to Ubiquitin 1; a.k.a. NEDD8 for Neural Precursor Cell Expressed Developmentally Down-Regulated 8 in mammals) are limited to one family of proteins, Cullins (7-10), although hundreds of targets can be Neddylated in human cells under certain conditions (11). Moreover, only few dedicated proteins have been identified to distinguish between Ub and Rub1, or dynamically regulate processing of Rub1 signals (12-15). The paucity of Rub1 targets and the divergence of Rub1 and Ub cellular landscapes are puzzling, given that the two proteins share ∼60% sequence identity, an identical tertiary fold (16), and key recognition elements on their surface (**Fig 1 A-C**). We note that genes encoding for Ub and Rub1 are found in all eukaryotes (17,18), although conservation of Ub is far greater than that of Rub1 orthologs (**Fig 1B**). Accordingly, throughout this text we retain the species-specific nomenclature for *RUB1/NEDD8* unless addressing general properties of the proteins, in which case we will refer to “Rub1”. Cullins are the best studied targets of Rub1/NEDD8 (in *S. cerevisiae* there are three cullins: yCul1/Cdc53, yCul3, and Rtt101, whereas mammals express seven cullin proteins; (7,19,20)). Conjugation of Rub1 to a specific lysine on cullins is mediated by the consecutive action of an E1 activating enzyme (Uba3-Ula1 dimer in *S. cerevisiae*; also referred to as NEDD8 activating enzyme, NAE), an E2 conjugating enzyme (Ubc12) and an E3 ligase (Rbx1) (19,21,22). Cullin neddylation is reversed by proteolytic activity of the CSN complex (a.k.a. COP9 Signalosome) (12,23-32). A number of seemingly unrelated proteins were also shown to be neddylated in different organisms (11,14,33-41), which appear to be reversed in some organisms by another deconjugase, SENP8/DEN1 (12,42). So far, the biological outcomes of conjugation of these non-cullin targets is unclear, nor is it clear to what extent enzymatic cascades responsible for “low level neddylation of non-cullin targets” and those responsible for modification of cullins overlap. Nevertheless, even considering “atypical neddylation”, the number of *bona-fide* ubiquitination targets is much greater, a direct outcome of an expanded repertoire of ubiquitination enzymes, primarily multiple E2s and myriad E3s (6). Moreover, a large portion of ubiquitination is polymeric via isopeptide linkages between surface amines (primarily lysine residues) on a proximal Ub unit (M1, K6, K11, K27, K29, K33, K48, K63) to the carboxyl-terminus of the distal unit (43-45). The most abundant linkages in polyUb are via lysine-48 (hereafter K48), a constitutive signal for targeting conjugated substrates for proteolysis by 26S proteasomes, and via K63, which generally increase upon stress (5,44-48). The result is a characteristic heterogeneous landscape of polyUb conjugates that contrasts a more limited repertoire of Rub1 modifications under steady state conditions (**Fig 1D**).

**Figure 1.**
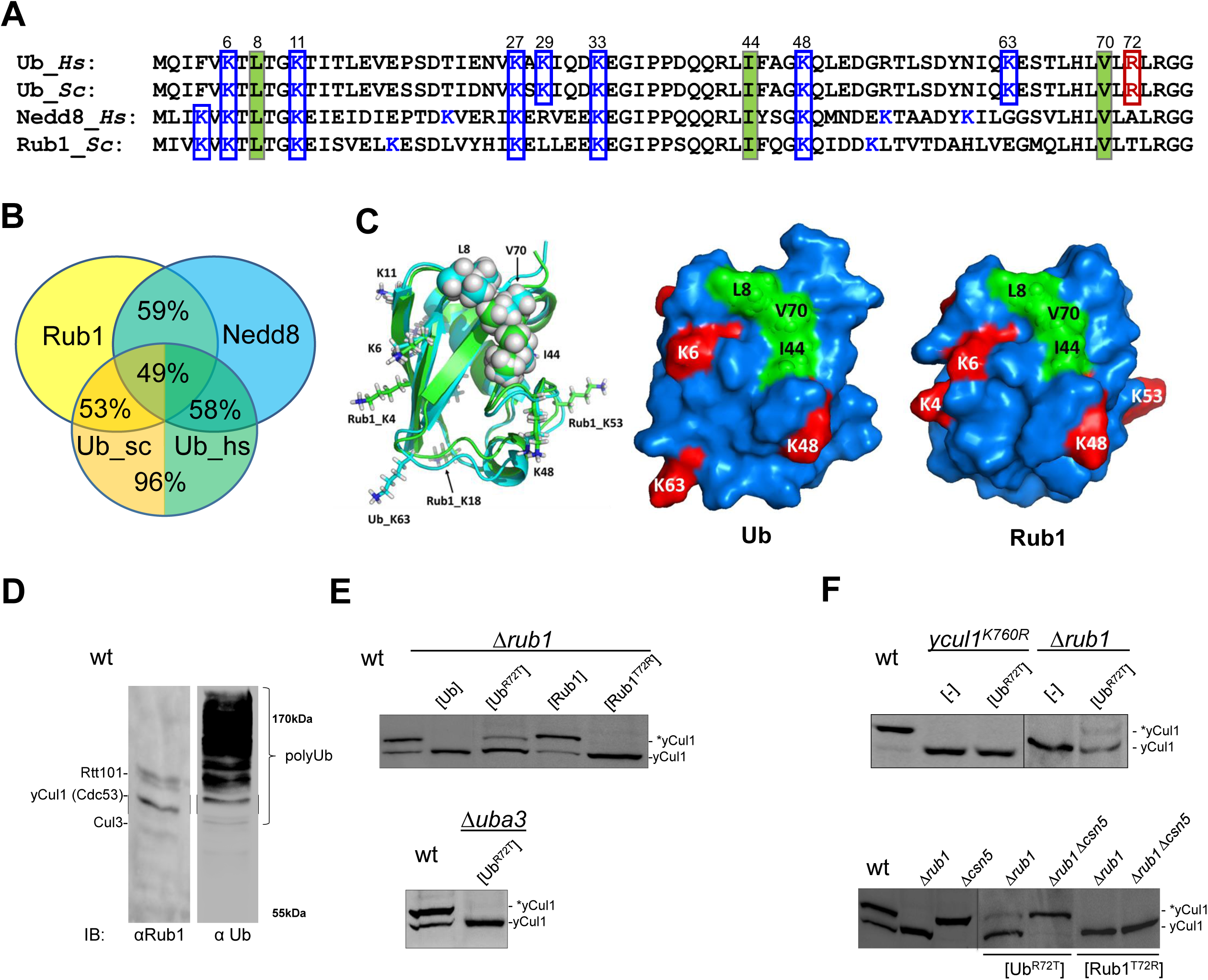
Ubiquitin R72T modifies yCul1, a neddylation target. (A) Clustal W multiple sequence alignment of Ub and Rub1/Nedd8 from *Saccharomyces cerevisiae* (Sc) and human (Hs). The hydrophobic patch residues of Ub (L8, I44, and V70) are framed in green. lysine residues are colored blue, and boxed if conserved in Ub or in Rub1. arginine 72 of Ub is colored red. (B) Percent sequence identity between Ub, ScRub1, and HsNEDD8. (C) Left: Structure alignment of Rub1 (cyan) and Ub (green). The hydrophobic patch residues are shown in sphere representation, the lysine side chains are shown as sticks. Consensus lysine side chains are indicated as sticks. Right: Surface representation of Ub and Rub1 from *Saccharomyces cerevisiae* based on published structures (16); surface hydrophobic patch is colored green, lysine residues are in red. (D) Whole cell extract of logarithmic *S. cerevisiae* culture resolved by 8% SDS PAGE and immuno-blotted for Rub1 (left) or Ub (right). Migration of neddylated yCul4/Rtt101, yCul1/Cdc53 and Cul3 is noted on left. (E, F) Whole cell extract of logarithmic phase wild-type, yCul1^K760/R^ or *rub1, csn5*, or *uba3* deletants and double mutants complemented with empty plasmids [-], or plasmids expressing proteins as noted, resolved by 8% SDS PAGE and immuno-blotted for yCul1/Cdc53. Lower band corresponds to unmodified yCul1. * marks migration of Rub1-yCul1.

E1 activating enzymes, the first enzymes in their respective modification cascades, distinguish between Ub and Rub1 despite their structural similarities. Thus, specificity of NAE for NEDD8/Rub1 is attributed to an arginine side chain (R190 in mammalian APPBP1-UBA3) that clashes with an arginine residue (R72) near the carboxyl tail of Ub if the latter enters the enzyme active site (49-52). Although the ubiquitin activating enzyme, UAE (a.k.a. UBA1) shows a preference for Ub over Rub1 (53), without a positive charge at position 72 of Rub1 (**Fig 1A**) some Rub1 is likely to engage UAE when presented at high levels (16,53-55). The leeway afforded by UAE to activate both Ub and Rub1 is one possible explanation for “atypical neddylation” of non-cullin targets upon overexpression of Rub1/NEDD8 or under stress that depletes levels of free Ub (16,54-57).

So far, the significance of “atypical neddylation” is unclear: is it an artifact of stressing the Rub1/NEDD8 modification pathway, or is it a natural manifestation of fluctuations in relative abundance of free Ub and Rub1/NEDD8? Since induction of NEDD8 or overreaction of the neddylation enzymatic cascade occurs in certain cancers, intensive therapeutic efforts have targeted NAE as a treatment (58). However, effects of NAE inhibitors appear to extend beyond their canonical role in modifying CRLs (59), emphasizing the need to fully map natural targets for NEDD8/Rub1 conjugation. In this regard, *S. cerevisiae* and *C. albicans* are potent model systems to test perturbations of the ubiquitin system, since they are among the very few organisms in which Rub1/NEDD8 is not essential (60-62). Depletion of Rub1 in these organisms would enable study of perturbations of both Rub1 and ubiquitin conjugation landscapes with only minimal perturbations to CRL-signaling pathways (since Cul1 is functional in *S. cerevisiae* even without Rub1). By designing reciprocal substitutions of residues 72 in both Rub1 and Ub, we directed Rub1 to modify targets of ubiquitination, and targeted ubiquitin to conjugate cullins. Crossover of Ub and Rub1 disrupted their cognate cascades without the inevitable perturbation that overexpression of a wild-type protein would have on its own pathway.

## RESULTS

### Replacement of Rub1 with ubiquitin in vivo

Even in the total absence of Rub1, Ub in *S. cerevisiae* did not spuriously modify yCul1 (**Fig 1E**), pointing to the stringent specificity of cullin-modification enzymes for Rub1. However, elimination of a single positive charge at position 72 of Ub (Ub^R72T^) led to Ub-modified cullins in cells (**Fig 1E**). Ub^R72T^ was activated by the E1 for Rub1, Uba3, and conjugated to yCul1 at the canonical neddylation site; i.e. K760 (**Fig 1E, F**), indicating it was recognized as Rub1 by key factors of the Rub1 modification cascade. An *in vitro* MS-based labeling method (63) confirmed that Ub^R72T^ was activated by NAE as efficiently as Rub1 (**Fig S1**). Ub-yCul1 conjugates were sensitive to hydrolysis by the CSN **(Fig 1F**), indicating that CSN recognizes modified cullins regardless of whether they are modified by Rub1 or Ub. Consistent with these observations, merely introducing a positive charge at position 72 of Rub1 (Rub1^T72R^) in cells eliminated modification of yCul1 (**Fig 1E)**. In a similar result, Rub1^T72R^ was not activated by NAE **(Fig S1)**, supporting the conclusion that NAE efficiently discriminates between Rub1 and ubiquitin. Apparently, a single amino acid is sufficient for E1s to distinguish between Ub and Rub1 in living cells at least in yeast and fungus and determine which E1 activates them and by extension, which targets are modified by Ub and which by Rub1. Thus, Ub^R72T^ was able to enter the Rub1 conjugation pathway and behave essentially as a “rubylized Ub”, a tool we then used to dissect the modification characteristics of Rub1 and Ub.

### Phenotypes of ubiquitin modifying Rub1 targets

With the information that Ub^R72T^ modifies typical targets of neddylation, we evaluated the fate of cullin modified by Ub instead of Rub1. Comparison of biological half-life demonstrated that Ub-modified yCul1 was turned over faster than unmodified or Rub1-modified yCul1 (**Fig 2A**). Turnover of Ub-modified yCul1 was likely proteasome-dependent since it was stabilized by treatment with a proteasome inhibitor, MG132 (**Fig 2B**). In order to substantiate the link between ubiquitination and proteolysis of cullin, we repeated the experiment with lysineless (K0) Ub that restricts modification of targets to a single (mono) Ub (44). Indeed, yCul1 modified by non-polymerizable Ub^K0,R72T^ was stable with no appreciable biological turnover (**Fig 2B**). Under steady-state conditions, the portion of Cul1 modified by Ub^K0,R72T^ out of total Cul1 in whole cell extract was comparable to the proportion of Rub1-modified Cul1 in the isogenic wild-type strain. In both cases, the steady state levels of modified cullin were greater than steady state levels of cullin modified with polymerizable Ub^R72T^ (**Fig 2C**). In absence of Csn5, the proteolytically active subunit of the CSN complex, Ub-modification of cullin – whether lysineless or polymerizable – increased (**Fig 2C; right**), supporting the observation from Fig 1F that CSN is not limited to Rub1 but rather is able to act on Cullin and deconjugate either Ub or Rub1.

**Figure 2.**
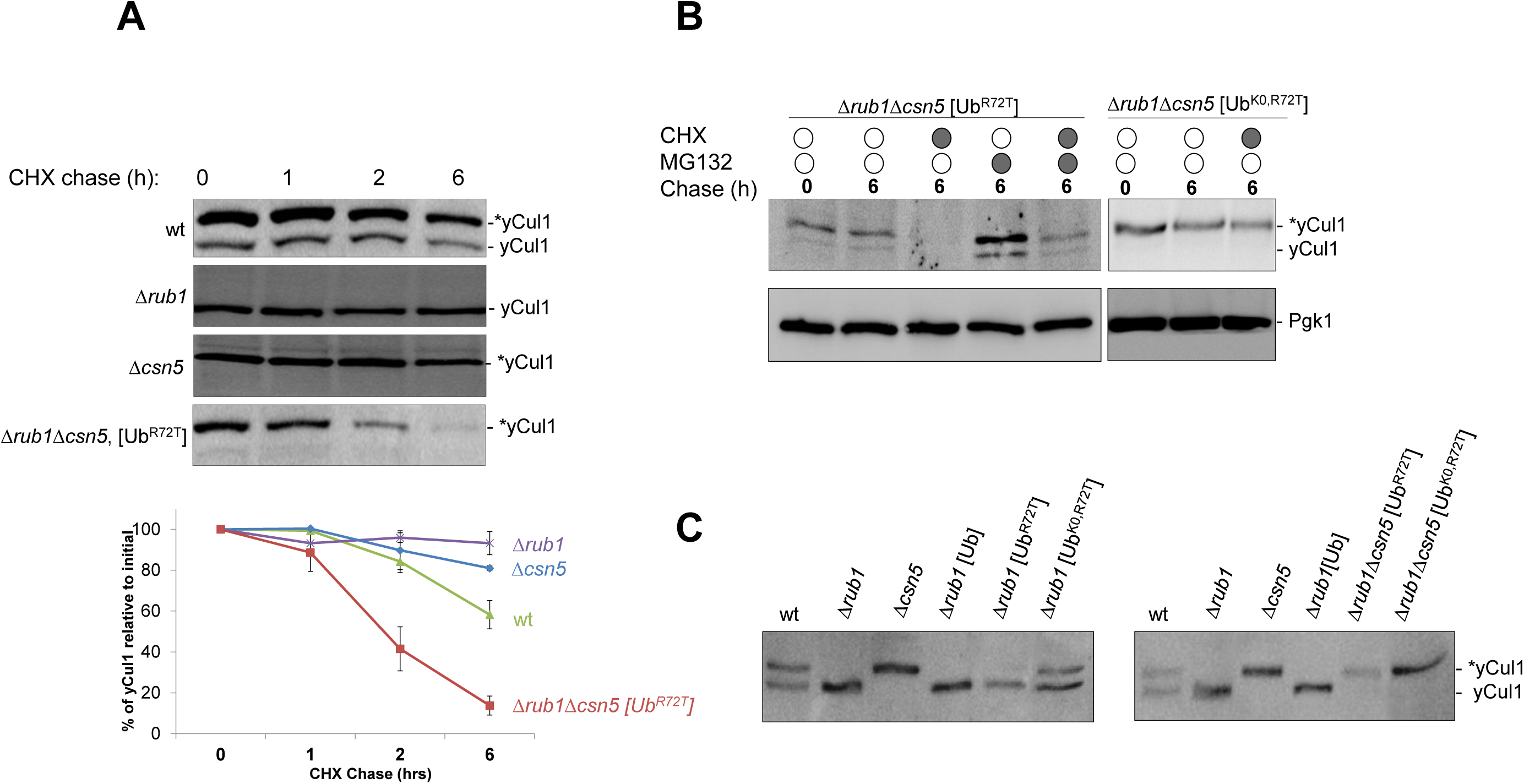
Ubiquitin-modified yCul1 is turned over faster than unmodified or Rub1-modified yCul1. (A) Half-life of Cul1. *S. cerevisiae* yeast strains as noted were grown to mid-log growth (t=0) and treated with 250µg/ml cyclohexamide (CHX). Cells were harvested at indicated times, resolved by 8% SDS PAGE and immuno-blotted for yCul1 (top). Quantification of immunoblots was performed using ImageJ v1.40f software (http://imagej.net/), relative intensity of 2 or more repeats was normalized to pre-treated yCul1 levels (t-0), and plotted over time (bottom). (B) Cell cultures of Δ*rub1*Δ*csn5* expressing either [Ub^R72T^] or [Ub^K0,R72T^] treated with 250 µg/ml CHX for 15 min before the addition of 75µM of proteasome inhibitor MG132 for 6 hours. Samples were resolved by 8% SDS PAGE and immunoblotted for yCul1 or Pgk1 as a loading control. (C) Whole cell extract of isogenic wild-type, Δ*rub1*, Δ*csn5* or double mutants expressing Ub mutants as noted, resolved by 8% SDS PAGE and immunoblotted for yCul1. * marks migration of Ub/Rub1-modified yCul1.

In order to substantiate the observation that cullins behave similarly when modified by either Rub1 or a non-polymerizable Ub, we checked whether non-polymerizable Ub could rescue phenotypes associated with loss of Rub1. In *S cerevisiae, Δrub1* phenotypes are mild, but include an effect on mitochondrial respiration (64). Indeed, the mild mitochondrial defects of *Δrub1* were reversed upon expression of a lysineless “rubylized” Ub: Ub^K0,R72T^ (**Fig S2**). Interestingly, lysine residues responsible for the three most prevalent linkages in polyubiquitin (polyUb) chains (K63, K48, and K11; (5,44)) are absent from the Rub1 sequence in *C. albicans* (**Fig S3)**, providing a unique opportunity to test the requirement for these lysine residues on the surface of Rub1. Deletion of *RUB1* in *C. albicans* caused a switch from yeast morphology in WT to hyphal growth in *rub1*^-/-^ (62), probably due to impairment of SCF E3 ligase complex activity causing increased switching from yeast to hyphal growth (65-67). Expression of either Ub^R72T^ and Ub^K0,R72T^in *rub1*^-/-^ reversed the *C. albicans* null hyphal growth phenotype (**Fig 3**). That non-polymerizable Ub rescues phenotypes of *rub1* null in two separate species lends support to the possibility that Rub1 does not require modifications on key surface lysines (including K48), at least as far as cullin modifications are concerned.

**Figure 3.**
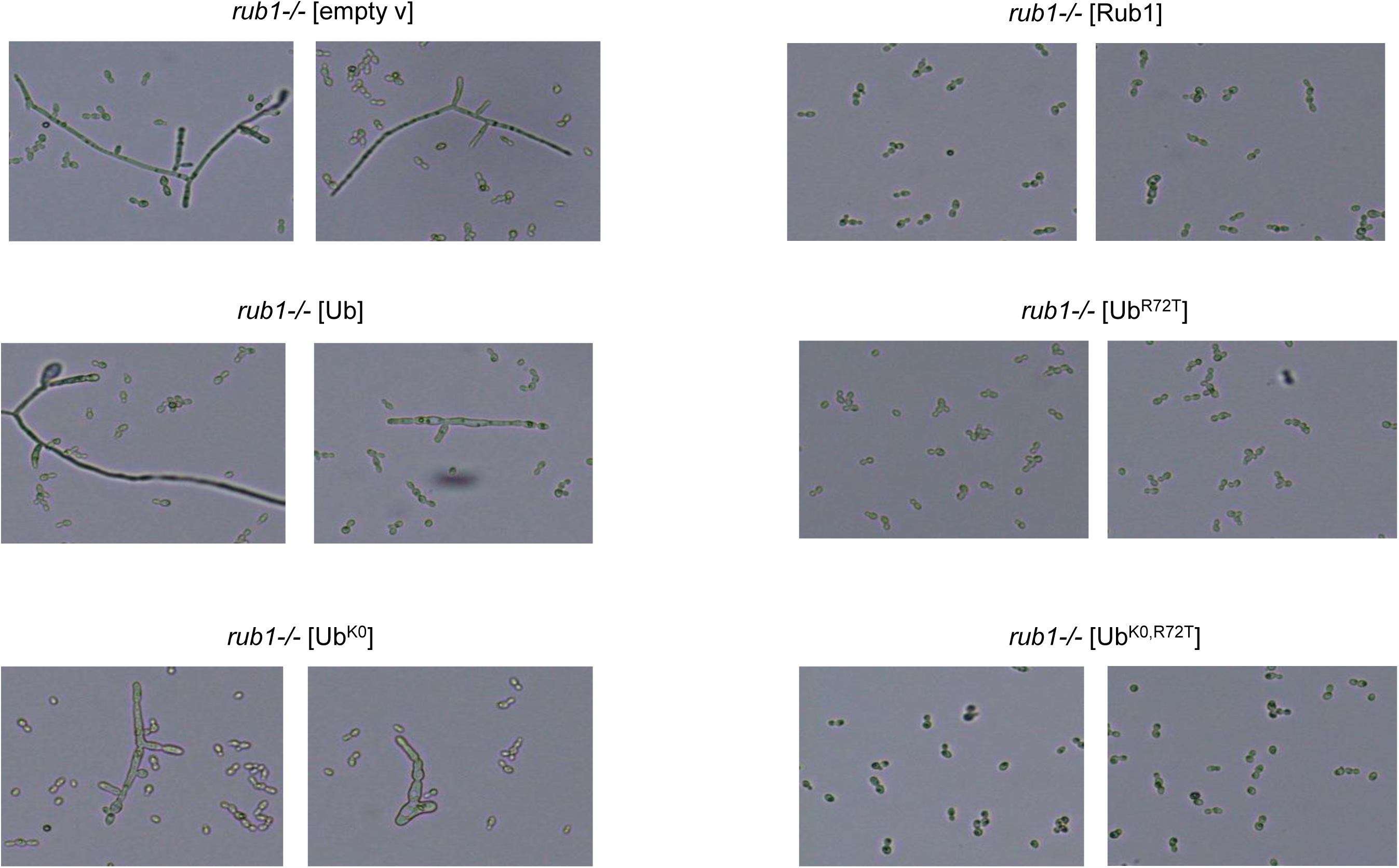
Ub^K0,R72T^ rescues Δ*rub1* phenotypes. *C. albicans* cells deleted for *rub1 (*Ca*rub1-/-)* and expressing Rub1 or Ub or mutated forms of Ub (Ub^K0^, Ub^R72T^ or Ub^K0,R72T^). Alterations in cell morphology were visualized by light microscopy.

### Mixed Rub1-Ubiquitin modifications

Having evaluated the effect of replacing Rub1 with Ub on cullins, we next assessed the outcome of Rub1 that is activated by the ubiquitin cascade, presumably modifying typical targets of ubiquitination. Overexpression of Rub1 led to accumulation of Rub1 in high MW conjugates (**Fig 4A**). The resulting heterogeneous profile of Rub1-conjugates resembles previous results showing an increased repertoire of NEDD8/Rub1 targets upon overexpression, many of which were also typical substrates for ubiquitination (13,16,55-57). No measurable decrease of these high MW Rub1-conjugates was observed in a *UBA3* deletion (**Fig S4**), indicating that formation of these conjugates was not performed by NAE but likely carried out by enzymes of the ubiquitination cascade. This phenomenon appears to be a common feature of both yeast and human cells. For instance, overexpression of NEDD8 in H1299 cells resulted in neddyylation of known ubiquitination targets, such as the HECT E3 ubiquitin ligase HERC2, RING ubiquitin E3 ligases Rlim, Livin-α, or the tumor suppressor p53 (**Fig S5A, B**). All four of these proteins can be highly ubiquitinated under certain conditions and subsequently degraded by proteasome (68-77). Moreover, since overexpression of MDM2, a well-documented E3 ubiquitin ligase for p53, significantly increased the extent of NEDD8-p53 conjugates (**Fig S5B, C**) it is likely that neddyylation of these targets also depends on enzymes of the ubiquitination cascade. Mdm2-mediated p53 neddylation has been reported (39), though whether this and other examples of atypical neddylation (11,13) reflect a dedicated pathway for NEDD8 modifications or a secondary role of ubiquitination enzymes has not been clarified for each case yet.

**Figure 4.**
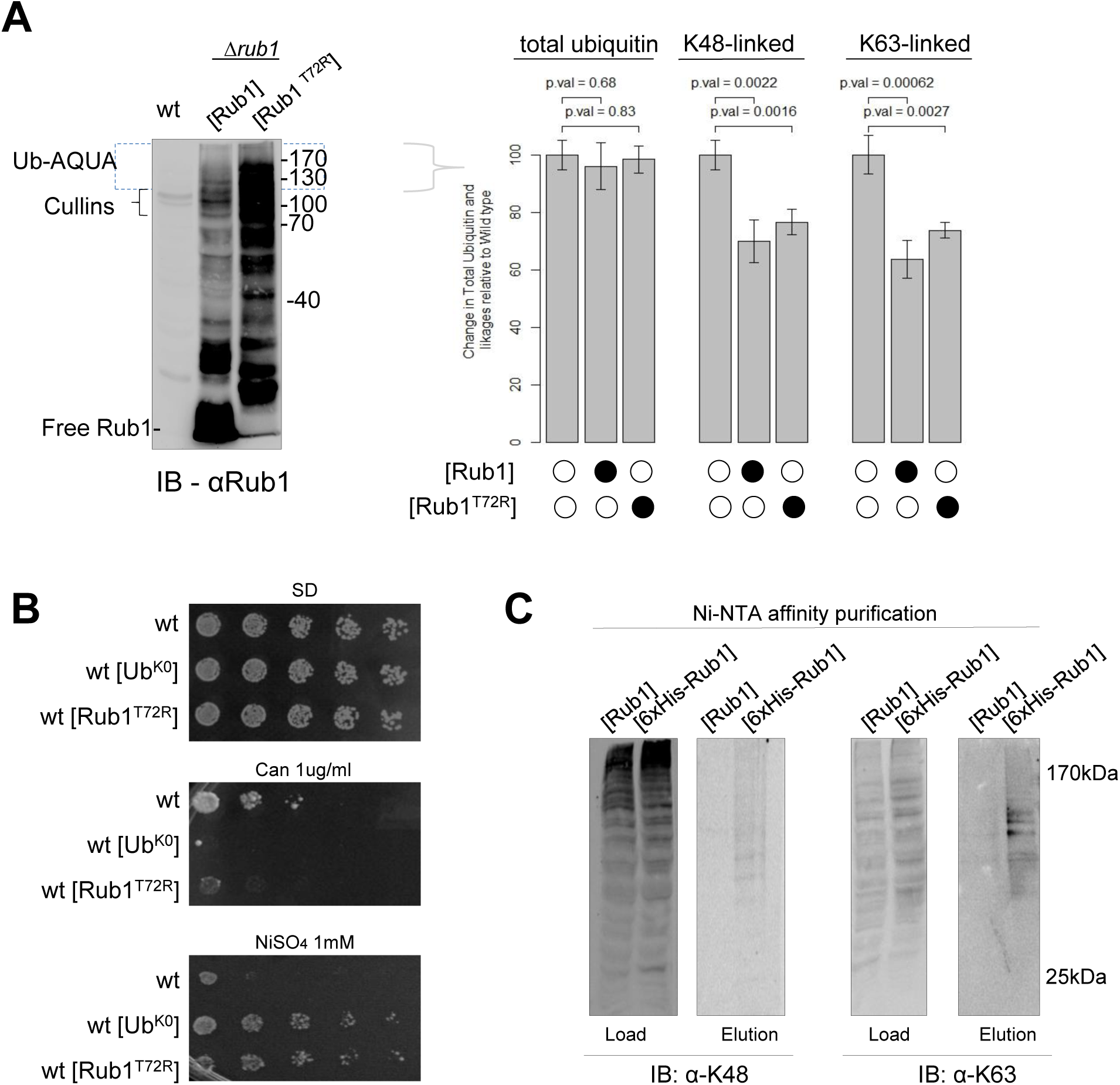
Effects of elevated Rub1 on the Ubiquitin landscape. (A) Right: Whole cell extracts from log phase wild-type, or wild-type cells overexpressing either Rub1 or Rub1^T72R^ resolved by 15% SDS PAGE and immuno-blotted for Rub1. Left: Top section of each of the lanes, corresponding to migration of proteins with molecular weight greater than ∼120 KDa (to exclude cullins) was excised, trypsinized and subject to quantitative MS/MS by Ub-AQUA following earlier protocols (44,109-112). K48- and K63-signature polyUb chain peptides were quantitated by comparison of signal intensity of isotopically labelled standards using mass spectrometry. Graph shows fold-change of polyUb chain linkages found in wild-type vs induced Rub1 cells. *Error bars* indicate standard deviation of three technical replicates. (B) Drop assay of wild-type or wild-type cells overexpressing either Ub^K0^ or Rub1^T72R^ were seeded in serial dilutions onto SD-agar plates (top) or SD-agar plates supplemented with the amino acid analog canavanine (middle) or nickel sulfate (bottom). (C) Whole cell extracts of Δ*rub1* cells overexpressing either Rub1 or His_6_-Rub1 were affinity purified by Ni-NTA chromatography. Samples of input (load) and imidazole elutions were resolved by SDA PAGE and immunoblotted with antibodies specific for K48-linked polyUb (left), or K63-linked polyUb (right).

One limitation of Rub1 overexpression is the difficulty to distinguish the effects of elevated Rub1 on typical targets of neddylation from the effects of simultaneous spillover of Rub1 into the ubiquitin landscape. Realizing that a single positive charge near the carboxyl tail of Ub is sufficient to block its activation by NAE (49-52), we designed a “ubiquitinized form of Rub1” to study the outcome of Rub1 on the ubiquitin system, without the inevitable perturbation that overexpression of Rub1 would have on NAE function. In cells, expression of Rub1^T72R^ resulted in accumulation of high MW conjugates of Rub1 accompanied with an almost complete depletion of free (unconjugated) Rub1 (**Fig 4A**). Deletion of *UBA3* had little discernable effect on the Rub1-conjugation pattern (with the exception of bands corresponding to modified cullins; **Fig S4**). Although Rub1^T72R^ was not activated by NAE (**Fig 1E; Fig S1**), it was activated by UAE *in vitro* as efficiently as Ub was (**Fig S6**). This propensity of UAE to activate Rub1^T72R^ is a likely explanation for how Rub1^T72R^ accumulated in high MW conjugates to a greater extent than similarly expressed Rub1 (**Fig 4A**). These data are also consistent with the reduced efficiency of UAE to activate Ub^R72K^ or Ub^R72A^ shown earlier (63). Introduction of ubiquitinized Rub1 could be a useful tool to probe the Ub system without greatly affecting the Rub1 signaling system because it is inert to NAE.

As Rub1 lacks lysine at position 63 (**Fig 1A**), it would be unable to form polymers with an equivalent conformation to polyUb K63-linked chains. Therefore, we hypothesized that introducing Rub1 into the ubiquitination cascade could alter the polyUb-linkage profile. Overexpression of *RUB1* resulted in a mild decrease of conjugated polyUb, particularly K63-linkages (**Fig S7**), suggesting that the two conjugation pathways affect each other. Using Ub AQUA, we quantified linkage types in high MW Ub-conjugates and found that overexpression of *RUB1* lead to a significant decrease of both K63- and K48-linkages in whole cell extract (**Fig 4A**, right). This result is remarkably similar to the outcome of expressing non-polymerizable lysineless Ub (Ub^K0^) in a WT background (44). Despite the potential to interfere with any of the seven lysine-linkage types, published phenotypes of Ub^K0^ expression related primarily to endocytosis and protein sorting that are largely driven by K63-linked polyUb chains (44,78). Likewise, Rub1 embedded into polyUb chains caused sensitivity to canavanine and conversely tolerance to nickel ions (**Fig 4B**), both hallmarks of defective protein trafficking documented for induction of non-polymerizable Ub (22,44,79-87). These phenotypes fit with attenuation of K63-Ub signaling. Evidence for simultaneous modifications by Rub1 and Ub on a single target was obtained by affinity purification of His_6_-Rub1 and staining for conjugated Ub (**Fig 4C**). The evidence for simultaneous modifications of a single target by Rub1 and Ub raises the possibility of mixed Ub/Rub1 modifications, either as mixed (heterologous) chains or alongside each other at different sites on the substrate. We note that both K48-linked and K63-linked polyUb chains co-purified with Rub1-modified substrates, albeit K63-linkages were relatively enriched. Together, these results suggest that Rub1 activated by ubiquitination enzymes directly influences the ubiquitin-linkage landscape.

### Rub1 incorporates into polyubiquitin chains upon heat stress

Having observed that the ratio of Rub1 to Ub may influence the nature of the conjugated ubiquitin landscape, we wished to consider conditions that may influence this ratio. Short-term heat stress increases polyubiquitin, leading to a drop in free unconjugated Ub levels (88). As free Ub is polymerized into high molecular weight (HMW) conjugates, the pool of free Ub may diminish, therefore we hypothesized that the resulting shortage of free Ub may facilitate encounter of Rub1 with the Ub E1 activating enzyme, UAE, effectively directing some Rub1 into the ubiquitin landscape. Indeed, following heat stress we observed a greater increase in polyUb chains in *Δrub1* relative to wild-type cells (**Fig 5A;** left). Conjugated Ub in HMW polyUb chains following short-term heat stress was up to three-fold greater in *Δrub1* compared to wild-type, as estimated by Ub AQUA, (**Fig 5A**). The effect was common to both K48- and K63-linkages, both of which increased in *Δrub1* to a greater extent than in wild-type (**Fig 5B**). This result suggests that presence of Rub1 in the ubiquitin landscape results in an apparent reduction of total conjugated ubiquitin. In order to evaluate the outcome of Rub1 that is activated by UAE and is thus potentially conjugated to targets of ubiquitination, we incubated “ubiquitinized” Rub1^T72R^ with UAE and ubiquitin E2 enzymes. We found that although lysine residues K48 and K11 are conserved between Ub and Rub1, two enzymes UBE2K and UBE2S that generate predominantly K48 and K11 linkages with ubiquitin, respectively, were able to attach Rub1 to Ub, yet had difficulties further modifying Rub1 (**Fig S8**). To summarize, it appears that when Rub1 is conjugated to ubiquitin, it acts as a general polyUb chain terminator, decreasing average length of polyUb modifications on targets. Even when activated by UAE and entering the ubiquitin landscape, in cases when Rub1 is attached directly to a substrate it would presumably be primarily a monomeric modification, in line with Rub1 serving primarily as a monomeric signal on its canonical targets.

**Figure 5.**
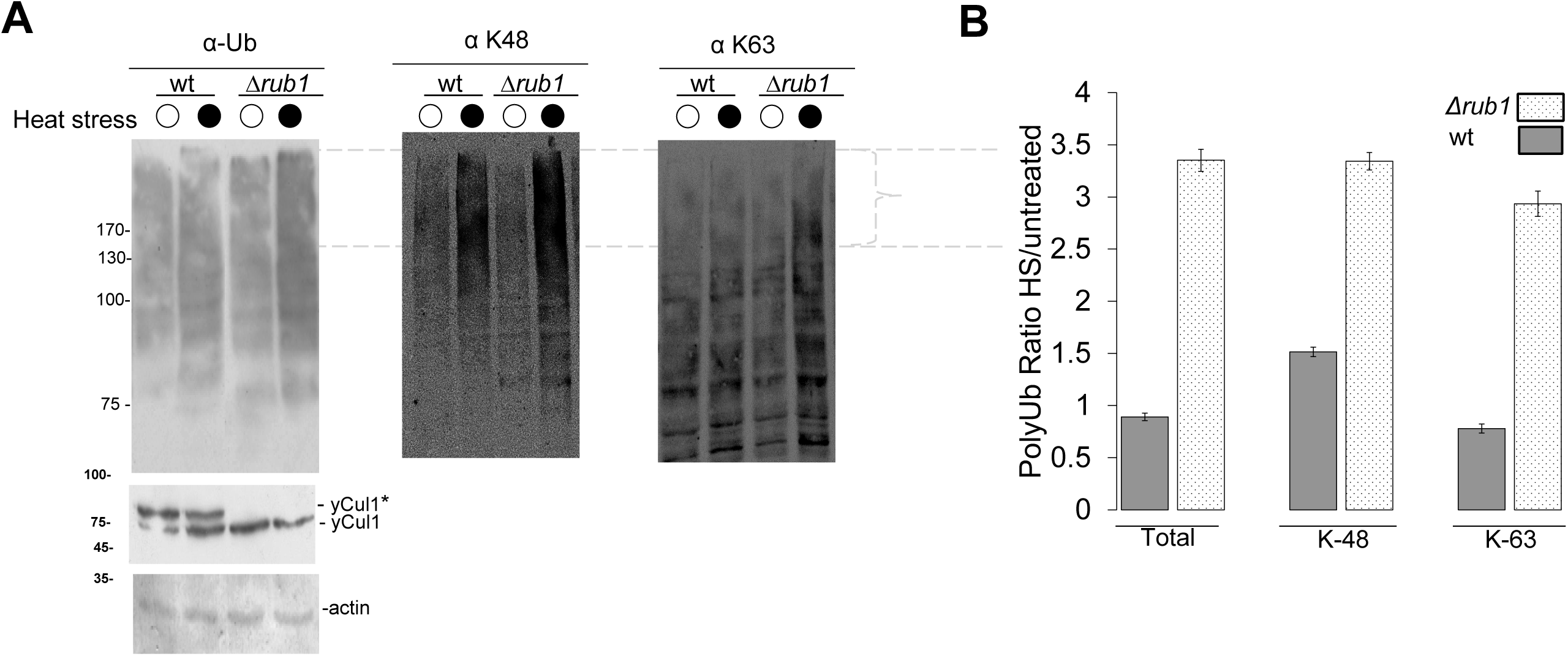
Abundance of polyUb chains increases in the absence of Rub1. (A) Cell cultures of wild-type or Δ*rub1* cells heat stressed for 30 minutes at 45°C were separated by SDS PAGE and blotted for ubiquitin, K48-Ub linkages or K63-Ub linkages using specific antibodies as indicated. Levels of yCul1 or actin serve as controls. (B) High molecular weight regions (above 100 kDa) were excised from gel and subjected for targeted mass spectrometry analysis for ubiquitin linkages types using Ub AQUA. Signature polyUb chain peptides were quantitated by comparison with isotopically labelled standards using mass spectrometry. Graph shows fold change of polyUb chain linkages found in wild-type or Δ*rub1* cell extracts. Error bars indicate standard deviation of three technical replicates of total Ub, K48- or K63-linkages measured in heat stress relative to untreated wild-type or Δ*rub1* cell extracts

## DISCUSSION

Although Ub and Rub1/NEDD8 are present as separate modifiers across Eukarya, this study suggests that they are partially interchangeable. Apparently, the cellular machinery allows for some naturally-occurring cross-activation, although we find greater leeway for one-directional cross-over of Rub1 into Ub signaling cascades rather than of Ub modifying typical targets of neddylation. There is mounting evidence for Rub1/NEDD8 conjugates beyond cullins. Targets of so-called atypical neddylation are prevalent, though when and to what purpose remains unclear. While Rub1 modifies some of these targets directly (10,11,89), the mere prevalence of Ub in the cell compounded by experimental difficulties in distinguishing between conjugated Rub1/NEDD8 and Ub (90,91) raises the possibility that a large portion of atypical neddylation is indirect through mixed chains. Overall, thousands of proteins may be modified by polymeric chains of ubiquitin, yet one of the most abundant cellular proteins, and one of the most heavily modified by ubiquitin, is Ub itself. This study touches on the outcome of Ub modified by Rub1, essentially forming mixed chains on targets.

By utilizing an experimental strategy that enhances cross-activation of Ub and Rub1, we addressed consequences of Ub or Rub1 entering each other’s respective signaling pathways, both in cells and in reconstituted enzymatic cascades. This experimental approach made it possible to study the perturbation that Rub1 introduces to ubiquitin signaling system without the inherent stress that overexpressed Rub1 would instill onto typical targets of neddylation. Apparently, residue 72 is the key to be recognized by E1 (either UAE or NAE), however, some ubiquitin E2 conjugating enzymes do recognize residues other than 72 and therefore Rub1 that does get activated by UAE is distributed differently than ubiquitin in the ubiquitin landscape. Moreover, some deubiquitinases also recognize unique surface of Ub and hence cleave Ub-Rub1 or Rub1-Ub conjugates differentially (92), in effect guaranteeing a distinct fate for mixed UBL-Ub polymers over polyubiquitin (although CSN or proteasome are able to remove Rub1 from Ub (16)). In this manner, introduction of Rub1 mutated at position 72 is a new tool for specifically affecting the Ub system without greatly affecting the Rub1 signaling system because it is inert to NAE. Thus ubiquitinized Rub1/NEDD8 may be added to other experimental tools for perturbing the ubiquitination landscape such as expression of lysineless non-polymerizable Ub (44), ubistatins (93,94), or the anti-cancer drug bortezomib (95).

Selection of either Ub or Rub1 by their preferred activating enzyme is largely dependent on the identity of residue 72 of the UBL (**Fig 6**). A single site substitution at position 72 was sufficient to enable NAE to activate Ub leading to cullin ubiquitination and subsequent degradation. UAE is naturally less selective than NAE, allowing some Rub1 to be activated by UAE. When the ratio of free Rub1 to Ub increases, as occurs under certain stress conditions such as heat stress or upon heterologous overexpression, greater is the chance that Rub1 could be activated by UAE. Such UAE-activated Rub1 is one mechanism that leads to “atypical Neddylation” of targets (**Fig 6**). Once activated by UAE, Rub1 can be relayed to E2 ubiquitin conjugating enzymes and eventually be conjugated to proteins, or to Ub generating mixed chains. Since E2 enzymes require information in the globular body of Ub beyond the C-terminal stretch required for activation by E1 (96) they may handle Rub1 differently than Ub and hence also participate in shaping the nature of the modification. Indeed, we found that some E2s were more efficient at attaching Rub1 to Ub than to Rub1, essentially terminating chain elongation with Rub1. This result is in line with Rub1 serving primarily as a monomeric modification on its typical substrates.

**Figure 6.**
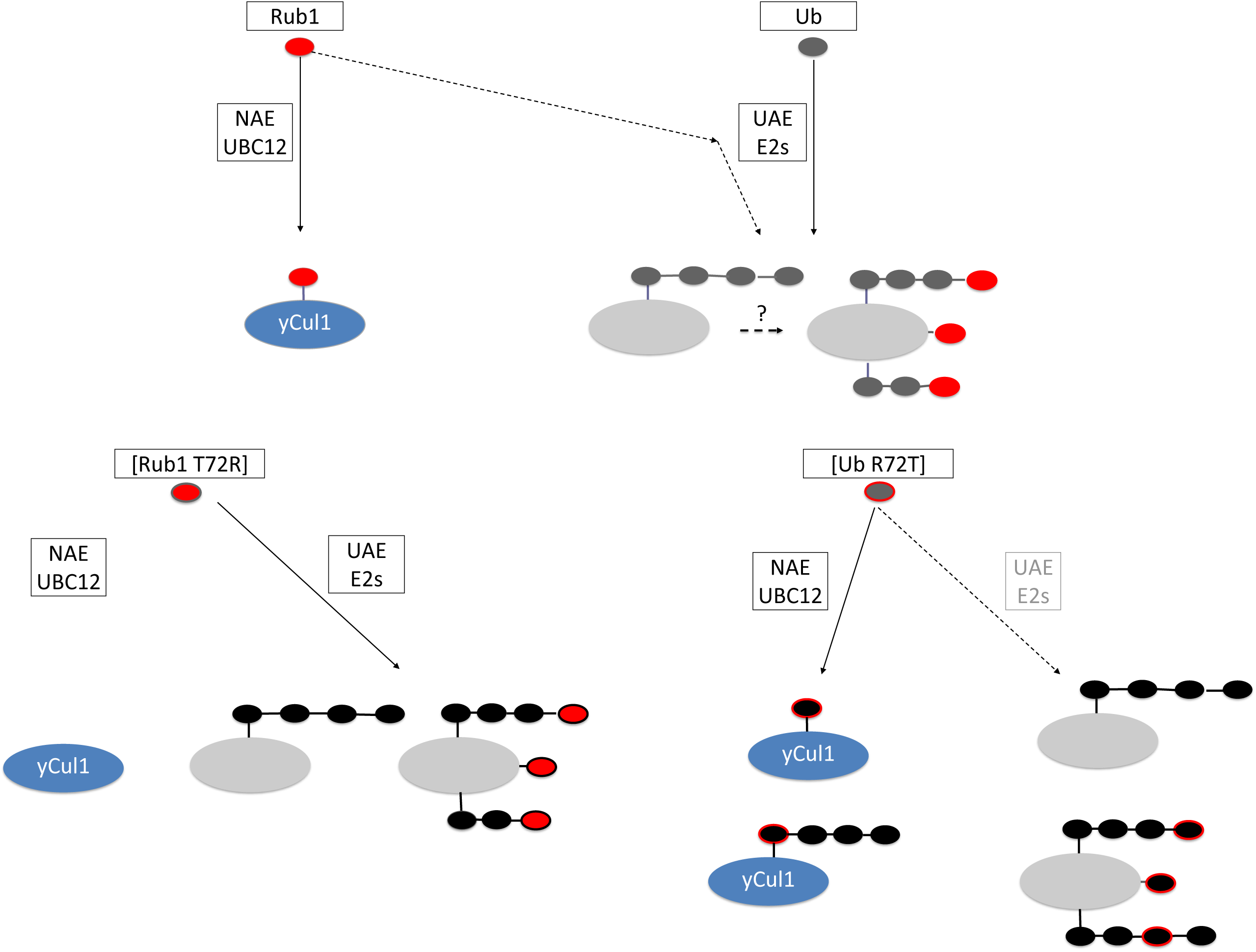
Model describing spillover of Rub1 into the ubiquitin system for protein modification. The Rub1/NEDD8 activating enzyme, NAE, stringently discriminates between NEDD8 and its close paralog Ub, showing a substantial preference for the former. By contrast, the Ub activating enzyme, UAE, is more flexible allowing some activation of Rub1 protein, which can account for some common targets of Ub and Rub1, including Ub itself. Ub E2 conjugating enzymes appear to be less efficient at modifying Rub1, hence the majority of Rub1 ends up as a terminal modification (as a monomeric signal or capping polymers of ubiquitin). Crossover of Rub1 into the Ub conjugating system can occur under certain stress conditions that alter the balance between available “free” forms of these small protein modifiers. A single-site mutation at position 72 of each protein generates proteins that retain all their properties as Ub/UBL signals yet are readily activated by enzymes of the respective pathways. The mutant proteins provide a powerful tool to probe the essence of the ubiquitin signal and the evolutionary constrains of these highly conserved protein signals.

When would cells need to regulate polyUb-chain length? Possibly, uncontrolled chain elongation is unnecessary or even detrimental for cells under normal growth conditions, especially given that four Ub units linked via K48 are an efficient proteasome-targeting signal (97,98). K63-linked polyUb chains are typically shorter (79,99,100). Short-term heat stress or acute oxidative stress are examples of conditions that entail greater need for protein removal and hence the need for enhanced ubiquitination activity. With a greater portion of Ub tied up in polyUb chains, a shortage of free Ub available for conjugation may ensue (101-103). As the ratio of free Ub to free Rub1 decreases, some Rub1 may be activated by UAE thereby entering the Ub landscape where it ends up modifying typical targets of ubiquitination. Our results suggest that introducing Rub1 into the ubiquitinome decreases the average length of heterologous mixed Rub1-Ub chains relative to homogenous polyUb, with a more pronounced effect on pathways that rely on K63-linked polyUb signaling.

As a small protein modifier, Rub1 seemingly “resembles monoUb” (rather than “<u>r</u>esembles <u>ub</u>iquitin”, as its name implies), explaining why some of its functions can be replaced by a lysineless non-polymerizable Ub (**Fig 2, 3**). The fundamental property of Ub as a polymeric signal may explain why NAE is more stringent in selecting Rub1 over Ub than UAE is in discriminating between the two. By preferring Rub1 to Ub, NAE essentially guarantees mono-modification of its targets with a single Ub-like domain (**Fig 6)**. Although Ub is the most conserved protein in eukaryotes (yeast and human orthologues differ by only three amino acids (104)), the protein is subject to variations in the form of post-translational modifications (PTMs; (47,105)). Ub itself is the most prevalent modifier of ubiquitin; roughly half of all Ub is modified on one or more lysine residues by another Ub molecule (5,44). The current study demonstrates that another Ub-like protein, Rub1/NEDD8, is also a Ub-modifier. One outcome of this modification is an interference in chain elongation. Since Rub1/NEDD8 lack a lysine residue at position 63, the direct outcome is on K63-linked chains, however we observed an effect on other linkages as well, probably due to the different ability of E2 enzymes to modify Rub1. We conclude that although the two proteins are quite similar (**Fig 1**), sharing a similar 3-D fold and key residues on their surface that can be recognized by similar receptors or processing enzymes (16,61,92), Rub1 and Ub diverge in their potential to polymerize or form elongated chains.

## EXPERIMENTAL PROCEDURES

### Strains, Plasmids and growth conditions

Strains and plasmids used in this study are described in Table S1 and Table S2. Transformations of yeasts with the relevant plasmids were performed by standard lithium acetate/polyethylene glycol 3350 procedure. Double mutants were produced through mating followed by sporulation and random spore analysis (106). Genotypes of haploid progeny were determined by mating with haploid reference strains, PCR and immunoblotting. Plasmids including mutated genes were constructed through PCR and were sequenced to confirm the positioning of the mutations. Maintenance of yeast strains was achieved by culturing plasmid-containing strains in a Synthetic Defined (SD) dropout medium composing of Yeast Nitrogen Base 0.69% [w/v] (Formedium, Hunstanton, Norfolk, UK), amino acid mix 0.14% [w/v] (Formedium), Dextrose 2%. Transformants were grown in SD medium excluding specific amino acids (leucine, tryptophan) or nucleotide (uracil). Experiments were performed when cells have reached an OD_600_ of 0.8. Yeast strains were grown at 30°C. For heat stress, cultures at 0.8 OD_600_ were incubated for 30 minutes at 45°C. For *Candida albicans*, an isogenic rub1-/- strain was grown and transformed as described (62).

### Drop assay

Cultures transformed by plasmids bearing either RGS-His_8_-K0 Ub or RGS-His_8_-Rub1^T72R^ were grown overnight. At the next morning, the cultures were harvested and washed twice with sterile distilled water. Finally, equal number of cells were confirmed by cell counting before five-fold serial dilutions in sterile double distilled water (DDW) followed by spotting 2 µl on SD agar plates with or without 1 µg/ml Canavanine, or 1 mM NiSO_4_. Plates were incubated at 30 °C.

### Microscopy

*Candida albicans* cells were visualized with a Zeiss AxioImager M1 microscope equipped with differential interference contrast (DIC) optics, using a 40× objective.

### Immunoblotting

Cells were harvested in trichloroacetic acid (TCA) as we previously described (107). After lysis, a buffering solution of 1 M Tris pH 11 was added to TCA lysed samples to neutralize any residual TCA. Laemmli buffer was added to all TCA precipitates before separation on SDS-PAGE. Experiments were repeated at least three times and a representative image is shown. In several cases, quantification of immunoblots was performed, using ImageJ v1.40f software (http://imagej.net/). Antibodies: Ubiqutin (Dakocytomation); Cdc53/yCul1 (Santa Cruz Biotechnology, Santa Cruz, CA sc-6717); K48-linked polyUb chains (Millipore 05-1307), K63-linked polyUb chains (Millipore 05-1308), Rub1/Nedd8 (Abcam ab4751-500). Actin (Abcam ab14128), Pgk1 (Abcam ab113687).

### Inhibition of protein synthesis and degradation

To follow after the decay of Cdc53/yCul1 expression in WT or mutant yeast cells, protein synthesis was inhibited by Cycloheximide (CHX). A freshly prepared 250 µg/ml CHX (from a stock of 10 mg/ml in DMSO) was added to equal amount of cultures for indicated times. To inhibit the turnover of Cdc53/yCul1, 75 µM of the proteasome inhibitor MG132 was added to the cultures, 15 minutes after adding CHX for 6 hours. Following the treatment, cells were harvested in TCA at indicated times and separated on an SDS-PAGE for immunoblotting.

### Isolation of neddylated conjugates

Isolation of conjugates was performed as described (44). In brief, WT cells transformed with either RGS-His_8_-WT Rub1 or untagged Rub1 at the logarithmic phase were lysed with glass beads in a 20% TCA solution. The final cell lysate of each sample was adjusted to 12% TCA for efficient protein precipitation. Cell lysates were incubated in ice for 1h, followed by full-speed centrifugation at 4C during 10min to separate the precipitated proteins from supernatants. The acidic pH of the precipitated proteins was adjusted to neutral using 1 M Tris pH11 and destined for nickel pullout were resuspended in loading buffer (6 M guanidine hydrochloride (GuHCl), 20 mM Tris pH8, 100 mM K_2_HPO_4_, 10 mM imidazole, 100 mM NaCl, 0.1% triton X-100). Equal amounts of each sample were loaded onto a mini NiNTA column (Qiagen, The Netherlands) and incubated at 4°C during 2 hours. The columns were subsequently washed with wash buffer 1 (20 mM Tris pH 8, 100 mM K_2_HPO_4_, 20 mM imidazole, 100 mM NaCl, 0.1% triton X-100), followed by washes with wash buffer 2 (20 mM Tris pH8, 100 mM K_2_HPO_4_, 10 mM imidazole, 1M NaCl, 0.1% triton X-100), followed by washes with wash buffer 3 (20 mM Tris pH8, 100 mM K_2_HPO_4_, 10 mM imidazole, 100 mM NaCl), followed by imidazole elution (500 mM imidazole). Elutions were concentrated using 12% TCA overnight at 4°C, followed by acetone precipitation at -20°C. The samples were then resuspended in Laemmli buffer and immunoblotted of K48- and K63-linked polyUb chains. Human NAE plasmid (pGST-E1-NEDD8) was obtained from Brenda Schulman and expressed in BL21 (DE3) cells and purified using GST affinity columns from GE.(108).

### Neddylation assays in mammalian cells

H1299 cells at 90% confluency in 6 cm plates were transfected using Lipofectamine 2000. After 20-24 hours transfection, the cells were lysed in 500 μl GuHCl buffer and 20 μl of protein A sepharose beads (equilibrated in guanidine hydrochloride, GuHCl buffer) were added. The samples were incubated at 4°C with rotation for 1 hour. After 1 hour, samples were centrifuged and 50 μl of Ni-Agarose beads (equilibrated in GuHCl buffer) were added to the supernatant followed by incubation at 4°C for 3-4 hours or overnight. The beads were then washed two times with GuHCl buffer followed by two more washes in a buffer containing one part of GuHCl buffer and four part of 50 mM Tris-Cl (pH 6.8) containing 20 mM imidazole. Finally, the samples were washed two times with 50 mM Tris-Cl (pH 6.8) buffer containing 20 mM imidazole. The samples were boiled at 95°C for 5 minutes in 100 μl of laemmli buffer containing 200 mM imidazole and loaded onto SDS-PAGE gels.

### Ubiquitin linkage quantification

We adopted a strategy to identify the linkage types of the polyUb chains that can be assembled through isopeptide bonds with any of the seven internal lysine residues. Each linkage type generates a unique peptide signature upon trypsin digestion that can be discriminated by mass spectrometry and quantitated by comparison with isotopically labelled standard. Whole cell extracts of yeast strains were resolved by 8% SDS PAGE. Top section of each lane corresponding to migration of proteins with molecular weight greater than ∼120 kDa (to exclude cullins) was excised. Proteins were digested in-gel using trypsin and subject to quantitative MS/MS by Ub-AQUA following earlier protocols (44,109-112). K48- and K63-signature polyUb chain peptides of were quantitated by comparison of signal intensity of isotopically labelled standards using mass spectrometry.

Samples from three technical replicates were loaded on an 8% stacking gel and proteins compressed in one band. Proteins were digested in-gel using trypsin, and purified as described above. The peptides were separated on a 25 cm reverse phase column (75 μm inner diameter, 3 μm reprosil beads, Dr. Maisch, GmbH, in house packed) using 5 to 50% acetonitrile gradient (VWR) with a flowrate of 250 nl/min (Eksigent nano-LC Ultra). The peptides were ionized on a Nano3 ion source (ABSciex) and directly sprayed into a QTRAP 5500 triple-quadrupole mass spectrometer, run in MRM mode to detect branched Ub-derived peptides and their synthetic isotopically labelled counterparts (44,113,114). Each sample was injected as technical triplicates. Two transitions for each peptide, i.e., UbTotal, Ub#14, UB#15, K48, K63, were acquired (supplementary Table S3). After the measurement peaks were integrated and light/heavy ratios calculated (115) using the Skyline software package (116) reflecting the abundance of the linkage type compared to the internal heavy standard.

Peptides ESTLHLVLR or TLSDYNIQK were used to measure the amount of Ub in the sample. The measurement of the two peptides were averaged. SRM conditions for the unlabeled ubiquitin peptides are listed in supplementary Table S3.

### Mass Spectrometry

All ESI-MS spectra were acquired on JEOL AccuTOF-CS mass spectrometer in a positive electrospray mode. High-resolution mass spectra of m/z 250-2500 were acquired for all samples. Spectra were deconvoluted using MagTran software with a 2-30 charge range combined with S/N of 3 to determine molecular mass.

## Acknowledgments

We would like to thank Oded Kleifeld for advice and suggestions on this work. Funding for this project was provided by an NSF-BSF grant MCB1818280 to MHG and DF, and Israel science foundation grants 2512/18 and 755/2019 to MHG and 162/17 to EP.

## Author contributions

SG performed most experiments in yeast and some in vitro assays. NR and SG performed molecular biology and genetic manipulations. AS, AT, NR and SG mapped ubiquitin landscape and Rub1 modifications. BEL performed most *in vitro* assays of Rub1 conjugation and synthesized mixed chains in vitro. RKS quantified E1 activity and carried out assays in mammalian cell culture. DK aided with experiments in *C albicans*. MS aided with experiments in mammalian cells. JL, OP, and GD performed mass spectrometry and Ub AQUA analysis. SG, NR, EP, DF, GD, and MHG designed experiments and monitored progress. All authors participated in writing and commenting on the manuscript. MHG, EP, DF, and GD wrote the first draft of the manuscript.

## Footnotes

i. Abbreviations used are: Ubiquitin-activating enzyme, UAE; Ubiquitin-activating enzyme, UBA; NEDD8-activating enzyme NAE; Ubiquitin conjugating enzyme, UBC; Cullin related ligase, CRL; Ubiquitin, Ub; Polyubiquitin, polyUb; Related to ubiquitin 1, Rub1; ubiquitin-like, UBL; Neural Precursor Cell Expressed Developmentally Down-Regulated 8, NEDD8; COP9 signalosome complex, CSN; guanidine hydrochloride, GuHCl
ii. in this manuscript we will preferentially refer to this ubiquitin-like modifier as Rub1 (to highlight its resemblance to ubiquitin), unless addressing specific properties of the mammalian gene product NEDD8.

## REFERENCES

1. Cappadocia, L., and Lima, C. D. (2018) Ubiquitin-like Protein Conjugation: Structures, Chemistry, and Mechanism. Chem Rev 118, 889–918

2. Taherbhoy, A. M., Schulman, B. A., and Kaiser, S. E. (2012) Ubiquitin-like modifiers. Essays Biochem 52, 51–63

3. Kerscher, O., Felberbaum, R., and Hochstrasser, M. (2006) Modification of proteins by ubiquitin and ubiquitin-like proteins. Annu Rev Cell Dev Biol 22, 159–180

4. Varshavsky, A. (2017) The Ubiquitin System, Autophagy, and Regulated Protein Degradation. Annu Rev Biochem 86, 123–128

5. Clague, M. J., Heride, C., and Urbe, S. (2015) The demographics of the ubiquitin system. Trends Cell Biol 25, 417–426

6. Glickman, M. H., and Ciechanover, A. (2002) The ubiquitin-proteasome proteolytic pathway: destruction for the sake of construction. Physiological reviews 82, 373–428

7. Deshaies, R. J., Emberley, E. D., and Saha, A. (2010) Control of cullin-ring ubiquitin ligase activity by nedd8. Sub-cellular biochemistry 54, 41–56

8. Rusnac, D. V., and Zheng, N. (2020) Structural Biology of CRL Ubiquitin Ligases. Advances in experimental medicine and biology 1217, 9–31

9. Wang, K., Deshaies, R. J., and Liu, X. (2020) Assembly and Regulation of CRL Ubiquitin Ligases. Advances in experimental medicine and biology 1217, 33–46

10. Baek, K., Krist, D. T., Prabu, J. R., Hill, S., Klugel, M., Neumaier, L. M., von Gronau, S., Kleiger, G., and Schulman, B. A. (2020) NEDD8 nucleates a multivalent cullin-RING-UBE2D ubiquitin ligation assembly. Nature 578, 461–466

11. Vogl, A. M., Phu, L., Becerra, R., Giusti, S. A., Verschueren, E., Hinkle, T. B., Bordenave, M. D., Adrian, M., Heidersbach, A., Yankilevich, P., Stefani, F. D., Wurst, W., Hoogenraad, C. C., Kirkpatrick, D. S., Refojo, D., and Sheng, M. (2020) Global site-specific neddylation profiling reveals that NEDDylated cofilin regulates actin dynamics. Nat Struct Mol Biol 27, 210–220

12. Schwechheimer, C. (2018) NEDD8-its role in the regulation of Cullin-RING ligases. Curr Opin Plant Biol 45, 112–119

13. Enchev, R. I., Schulman, B. A., and Peter, M. (2015) Protein neddylation: beyond cullin-RING ligases. Nat Rev Mol Cell Biol 16, 30–44

14. Xirodimas, D. P. (2008) Novel substrates and functions for the ubiquitin-like molecule NEDD8. Biochem Soc Trans 36, 802–806

15. Santonico, E. (2020) Old and New Concepts in Ubiquitin and NEDD8 Recognition. Biomolecules 10

16. Singh, R. K., Zerath, S., Kleifeld, O., Scheffner, M., Glickman, M. H., and Fushman, D. (2012) Recognition and cleavage of related to ubiquitin 1 (Rub1) and Rub1-ubiquitin chains by components of the ubiquitin-proteasome system. Mol Cell Proteomics 11, 1595–1611

17. Burroughs, A. M., Iyer, L. M., and Aravind, L. (2012) Structure and evolution of ubiquitin and ubiquitin-related domains. Methods Mol Biol 832, 15–63

18. Barth, E., Hubler, R., Baniahmad, A., and Marz, M. (2016) The Evolution of COP9 Signalosome in Unicellular and Multicellular Organisms. Genome Biol Evol 8, 1279–1289

19. Laplaza, J. M., Bostick, M., Scholes, D. T., Curcio, M. J., and Callis, J. (2004) Saccharomyces cerevisiae ubiquitin-like protein Rub1 conjugates to cullin proteins Rtt101 and Cul3 in vivo. Biochem J 377, 459–467

20. Willems, A. R., Schwab, M., and Tyers, M. (2004) A hitchhiker’s guide to the cullin ubiquitin ligases: SCF and its kin. Biochim Biophys Acta 1695, 133–170

21. Liakopoulos, D., Doenges, G., Matuschewski, K., and Jentsch, S. (1998) A novel protein modification pathway related to the ubiquitin system. EMBO J 17, 2208–2214

22. Finley, D., Ulrich, H. D., Sommer, T., and Kaiser, P. (2012) The ubiquitin-proteasome system of Saccharomyces cerevisiae. Genetics 192, 319–360

23. Pick, E., Hofmann, K., and Glickman, M. H. (2009) PCI complexes: Beyond the proteasome, CSN, and eIF3 Troika. Mol Cell 35, 260–264

24. Lingaraju, G. M., Bunker, R. D., Cavadini, S., Hess, D., Hassiepen, U., Renatus, M., Fischer, E. S., and Thoma, N. H. (2014) Crystal structure of the human COP9 signalosome. Nature 512, 161–165

25. Schmaler, T., and Dubiel, W. (2010) Control of Deneddylation by the COP9 Signalosome. Sub-cellular biochemistry 54, 57–68

26. Ambroggio, X. I., Rees, D. C., and Deshaies, R. J. (2004) JAMM: a metalloprotease-like zinc site in the proteasome and signalosome. PLoS Biol 2, E2

27. Cope, G. A., Suh, G. S., Aravind, L., Schwarz, S. E., Zipursky, S. L., Koonin, E. V., and Deshaies, R. J. (2002) Role of predicted metalloprotease motif of Jab1/Csn5 in cleavage of Nedd8 from Cul1. Science 298, 608–611

28. Enchev, R. I., Schreiber, A., Beuron, F., and Morris, E. P. (2010) Structural insights into the COP9 signalosome and its common architecture with the 26S proteasome lid and eIF3. Structure 18, 518–527

29. Maytal-Kivity, V., Reis, N., Hofmann, K., and Glickman, M. H. (2002) MPN+, a putative catalytic motif found in a subset of MPN domain proteins from eukaryotes and prokaryotes, is critical for Rpn11 function. BMC Biochem 3, 28

30. Birol, M., Enchev, R. I., Padilla, A., Stengel, F., Aebersold, R., Betzi, S., Yang, Y., Hoh, F., Peter, M., Dumas, C., and Echalier, A. (2014) Structural and biochemical characterization of the Cop9 signalosome CSN5/CSN6 heterodimer. PLoS One 9, e105688

31. Dubiel, D., Rockel, B., Naumann, M., and Dubiel, W. (2015) Diversity of COP9 signalosome structures and functional consequences. FEBS Lett 589, 2507–2513

32. Mosadeghi, R., Reichermeier, K. M., Winkler, M., Schreiber, A., Reitsma, J. M., Zhang, Y., Stengel, F., Cao, J., Kim, M., Sweredoski, M. J., Hess, S., Leitner, A., Aebersold, R., Peter, M., Deshaies, R. J., and Enchev, R. I. (2016) Structural and kinetic analysis of the COP9-Signalosome activation and the cullin-RING ubiquitin ligase deneddylation cycle. eLife 5

33. Jones, J., Wu, K., Yang, Y., Guerrero, C., Nillegoda, N., Pan, Z. Q., and Huang, L. (2008) A targeted proteomic analysis of the ubiquitin-like modifier nedd8 and associated proteins. J Proteome Res 7, 1274–1287

34. Loftus, S. J., Liu, G., Carr, S. M., Munro, S., and La Thangue, N. B. (2012) NEDDylation regulates E2F-1-dependent transcription. EMBO Rep 13, 811–818

35. Shu, J., Liu, C., Wei, R., Xie, P., He, S., and Zhang, L. (2016) Nedd8 targets ubiquitin ligase Smurf2 for neddylation and promote its degradation. Biochem Biophys Res Commun 474, 51–56

36. Xie, P., Zhang, M., He, S., Lu, K., Chen, Y., Xing, G., Lu, Y., Liu, P., Li, Y., Wang, S., Chai, N., Wu, J., Deng, H., Wang, H. R., Cao, Y., Zhao, F., Cui, Y., Wang, J., He, F., and Zhang, L. (2014) The covalent modifier Nedd8 is critical for the activation of Smurf1 ubiquitin ligase in tumorigenesis. Nat Commun 5, 3733

37. Mahata, B., Sundqvist, A., and Xirodimas, D. P. (2012) Recruitment of RPL11 at promoter sites of p53-regulated genes upon nucleolar stress through NEDD8 and in an Mdm2-dependent manner. Oncogene 31, 3060–3071

38. Sundqvist, A., Liu, G., Mirsaliotis, A., and Xirodimas, D. P. (2009) Regulation of nucleolar signalling to p53 through NEDDylation of L11. EMBO Rep 10, 1132–1139

39. Xirodimas, D. P., Saville, M. K., Bourdon, J. C., Hay, R. T., and Lane, D. P. (2004) Mdm2-mediated NEDD8 conjugation of p53 inhibits its transcriptional activity. Cell 118, 83–97

40. Xirodimas, D. P., Sundqvist, A., Nakamura, A., Shen, L., Botting, C., and Hay, R. T. (2008) Ribosomal proteins are targets for the NEDD8 pathway. EMBO Rep 9, 280–286

41. Hakenjos, J. P., Bejai, S., Ranftl, Q., Behringer, C., Vlot, A. C., Absmanner, B., Hammes, U., Heinzlmeir, S., Kuster, B., and Schwechheimer, C. (2013) ML3 is a NEDD8- and ubiquitin-modified protein. Plant Physiol 163, 135–149

42. Coleman, K. E., Bekes, M., Chapman, J. R., Crist, S. B., Jones, M. J., Ueberheide, B. M., and Huang, T. T. (2017) SENP8 limits aberrant neddylation of NEDD8 pathway components to promote cullin-RING ubiquitin ligase function. eLife 6

43. Castaneda, C. A., Dixon, E., Chaturvedi, A., Krueger, S., Cropp, T. A., and Fushman, D. (2013) Effect of Different Lysine Linkages on Polyubiquitin Chain Structure and Function. Biophysical Journal 104, 19a–19a

44. Ziv, I., Matiuhin, Y., Kirkpatrick, D. S., Erpapazoglou, Z., Leon, S., Pantazopoulou, M., Kim, W., Gygi, S. P., Haguenauer-Tsapis, R., Reis, N., Glickman, M. H., and Kleifeld, O. (2011) A perturbed ubiquitin landscape distinguishes between ubiquitin in trafficking and in proteolysis. Mol Cell Proteomics 10, M111 009753

45. Akutsu, M., Dikic, I., and Bremm, A. (2016) Ubiquitin chain diversity at a glance. J Cell Sci 129, 875–880

46. Heride, C., Urbe, S., and Clague, M. J. (2014) Ubiquitin code assembly and disassembly. Curr Biol 24, R215–220

47. Ohtake, F., and Tsuchiya, H. (2017) The emerging complexity of ubiquitin architecture. J Biochem 161, 125–133

48. Swatek, K. N., and Komander, D. (2016) Ubiquitin modifications. Cell research 26, 399–422

49. Bohnsack, R. N., and Haas, A. L. (2003) Conservation in the mechanism of Nedd8 activation by the human AppBp1-Uba3 heterodimer. J Biol Chem 278, 26823–26830

50. Walden, H., Podgorski, M. S., Huang, D. T., Miller, D. W., Howard, R. J., Minor, D. L., Jr., Holton, J. M., and Schulman, B. A. (2003) The structure of the APPBP1-UBA3-NEDD8-ATP complex reveals the basis for selective ubiquitin-like protein activation by an E1. Mol Cell 12, 1427–1437

51. Walden, H., Podgorski, M. S., and Schulman, B. A. (2003) Insights into the ubiquitin transfer cascade from the structure of the activating enzyme for NEDD8. Nature 422, 330–334

52. Souphron, J., Waddell, M. B., Paydar, A., Tokgoz-Gromley, Z., Roussel, M. F., and Schulman, B. A. (2008) Structural dissection of a gating mechanism preventing misactivation of ubiquitin by NEDD8’s E1. Biochemistry 47, 8961–8969

53. Whitby, F. G., Xia, G., Pickart, C. M., and Hill, C. P. (1998) Crystal structure of the human ubiquitin-like protein NEDD8 and interactions with ubiquitin pathway enzymes. J Biol Chem 273, 34983–34991

54. Hjerpe, R., Thomas, Y., Chen, J., Zemla, A., Curran, S., Shpiro, N., Dick, L. R., and Kurz, T. (2012) Changes in the ratio of free NEDD8 to ubiquitin triggers NEDDylation by ubiquitin enzymes. Biochem J 441, 927–936

55. Hjerpe, R., Thomas, Y., and Kurz, T. (2012) NEDD8 overexpression results in neddylation of ubiquitin substrates by the ubiquitin pathway. J Mol Biol 421, 27–29

56. Leidecker, O., Matic, I., Mahata, B., Pion, E., and Xirodimas, D. P. (2012) The ubiquitin E1 enzyme Ube1 mediates NEDD8 activation under diverse stress conditions. Cell Cycle 11, 1142–1150

57. Maghames, C. M., Lobato-Gil, S., Perrin, A., Trauchessec, H., Rodriguez, M. S., Urbach, S., Marin, P., and Xirodimas, D. P. (2018) NEDDylation promotes nuclear protein aggregation and protects the Ubiquitin Proteasome System upon proteotoxic stress. Nat Commun 9, 4376

58. Gatel, P., Piechaczyk, M., and Bossis, G. (2020) Ubiquitin, SUMO, and Nedd8 as Therapeutic Targets in Cancer. Advances in experimental medicine and biology 1233, 29–54

59. Mao, H., and Sun, Y. (2020) Neddylation-Independent Activities of MLN4924. Advances in experimental medicine and biology 1217, 363–372

60. Maytal-Kivity, V., Piran, R., Pick, E., Hofmann, K., and Glickman, M. H. (2002) COP9 signalosome components play a role in the mating pheromone response of S. cerevisiae. EMBO Rep 3, 1215–1221

61. Linghu, B., Callis, J., and Goebl, M. G. (2002) Rub1p processing by Yuh1p is required for wild-type levels of Rub1p conjugation to Cdc53p. Eukaryot Cell 1, 491–494

62. Sela, N., Atir-Lande, A., and Kornitzer, D. (2012) Neddylation and CAND1 independently stimulate SCF ubiquitin ligase activity in Candida albicans. Eukaryot Cell 11, 42–52

63. Singh, R. K., Kazansky, Y., Wathieu, D., and Fushman, D. (2017) Hydrophobic Patch of Ubiquitin is Important for its Optimal Activation by Ubiquitin Activating Enzyme E1. Anal Chem 89, 7852–7860

64. Bramasole, L., Sinha, A., Gurevich, S., Radzinski, M., Klein, Y., Panat, N., Gefen, E., Rinaldi, T., Jimenez-Morales, D., Johnson, J., Krogan, N. J., Reis, N., Reichmann, D., Glickman, M. H., and Pick, E. (2019) Proteasome lid bridges mitochondrial stress with Cdc53/Cullin1 NEDDylation status. Redox biology 20, 533–543

65. Atir-Lande, A., Gildor, T., and Kornitzer, D. (2005) Role for the SCFCDC4 ubiquitin ligase in Candida albicans morphogenesis. Mol Biol Cell 16, 2772–2785

66. Trunk, K., Gendron, P., Nantel, A., Lemieux, S., Roemer, T., and Raymond, M. (2009) Depletion of the cullin Cdc53p induces morphogenetic changes in Candida albicans. Eukaryot Cell 8, 756–767

67. Mendelsohn, S., Pinsky, M., Weissman, Z., and Kornitzer, D. (2017) Regulation of the Candida albicans Hypha-Inducing Transcription Factor Ume6 by the CDK1 Cyclins Cln3 and Hgc1. mSphere 2

68. Bach, I., Rodriguez-Esteban, C., Carriere, C., Bhushan, A., Krones, A., Rose, D. W., Glass, C. K., Andersen, B., Izpisua Belmonte, J. C., and Rosenfeld, M. G. (1999) RLIM inhibits functional activity of LIM homeodomain transcription factors via recruitment of the histone deacetylase complex. Nature genetics 22, 394–399

69. Garcia-Cano, J., Martinez-Martinez, A., Sala-Gaston, J., Pedrazza, L., and Rosa, J. L. (2019) HERCing: Structural and Functional Relevance of the Large HERC Ubiquitin Ligases. Front Physiol 10, 1014

70. Haupt, Y., Maya, R., Kazaz, A., and Oren, M. (1997) Mdm2 promotes the rapid degradation of p53. Nature 387, 296–299

71. Kasof, G. M., and Gomes, B. C. (2001) Livin, a novel inhibitor of apoptosis protein family member. J Biol Chem 276, 3238–3246

72. Kubbutat, M. H., Jones, S. N., and Vousden, K. H. (1997) Regulation of p53 stability by Mdm2. Nature 387, 299–303

73. Kuhnle, S., Kogel, U., Glockzin, S., Marquardt, A., Ciechanover, A., Matentzoglu, K., and Scheffner, M. (2011) Physical and functional interaction of the HECT ubiquitin-protein ligases E6AP and HERC2. J Biol Chem 286, 19410–19416

74. Ostendorff, H. P., Bossenz, M., Mincheva, A., Copeland, N. G., Gilbert, D. J., Jenkins, N. A., Lichter, P., and Bach, I. (2000) Functional characterization of the gene encoding RLIM, the corepressor of LIM homeodomain factors. Genomics 69, 120–130

75. Ostendorff, H. P., Peirano, R. I., Peters, M. A., Schluter, A., Bossenz, M., Scheffner, M., and Bach, I. (2002) Ubiquitination-dependent cofactor exchange on LIM homeodomain transcription factors. Nature 416, 99–103

76. Wu, W., Sato, K., Koike, A., Nishikawa, H., Koizumi, H., Venkitaraman, A. R., and Ohta, T. (2010) HERC2 is an E3 ligase that targets BRCA1 for degradation. Cancer Res 70, 6384–6392

77. Yang, Y., Fang, S., Jensen, J. P., Weissman, A. M., and Ashwell, J. D. (2000) Ubiquitin protein ligase activity of IAPs and their degradation in proteasomes in response to apoptotic stimuli. Science 288, 874–877

78. Hovsepian, J., Becuwe, M., Kleifeld, O., Glickman, M. H., and Leon, S. (2016) Studying Protein Ubiquitylation in Yeast. Methods Mol Biol 1449, 117–142

79. Erpapazoglou, Z., Dhaoui, M., Pantazopoulou, M., Giordano, F., Mari, M., Leon, S., Raposo, G., Reggiori, F., and Haguenauer-Tsapis, R. (2012) A dual role for K63-linked ubiquitin chains in multivesicular body biogenesis and cargo sorting. Mol Biol Cell 23, 2170–2183

80. Arita, A., Zhou, X., Ellen, T. P., Liu, X., Bai, J., Rooney, J. P., Kurtz, A., Klein, C. B., Dai, W., Begley, T. J., and Costa, M. (2009) A genome-wide deletion mutant screen identifies pathways affected by nickel sulfate in Saccharomyces cerevisiae. BMC Genomics 10, 524

81. Ruotolo, R., Marchini, G., and Ottonello, S. (2008) Membrane transporters and protein traffic networks differentially affecting metal tolerance: a genomic phenotyping study in yeast. Genome Biol 9, R67

82. Chen, Y., and Piper, P. W. (1995) Consequences of the overexpression of ubiquitin in yeast: elevated tolerances of osmostress, ethanol and canavanine, yet reduced tolerances of cadmium, arsenite and paromomycin. Biochim Biophys Acta 1268, 59–64

83. Grenson, M., Mousset, M., Wiame, J. M., and Bechet, J. (1966) Multiplicity of the amino acid permeases in Saccharomyces cerevisiae. I. Evidence for a specific arginine-transporting system. Biochim Biophys Acta 127, 325–338

84. Miura, T., and Abe, F. (2004) Multiple ubiquitin-specific protease genes are involved in degradation of yeast tryptophan permease Tat2 at high pressure. FEMS microbiology letters 239, 171–179

85. Becuwe, M., and Leon, S. (2014) Integrated control of transporter endocytosis and recycling by the arrestin-related protein Rod1 and the ubiquitin ligase Rsp5. eLife 3

86. Leon, S., and Haguenauer-Tsapis, R. (2009) Ubiquitin ligase adaptors: regulators of ubiquitylation and endocytosis of plasma membrane proteins. Exp Cell Res 315, 1574–1583

87. Nishimura, K., Igarashi, K., and Kakinuma, Y. (1998) Proton gradient-driven nickel uptake by vacuolar membrane vesicles of Saccharomyces cerevisiae. J Bacteriol 180, 1962–1964

88. Fang, N. N., Chan, G. T., Zhu, M., Comyn, S. A., Persaud, A., Deshaies, R. J., Rotin, D., Gsponer, J., and Mayor, T. (2014) Rsp5/Nedd4 is the main ubiquitin ligase that targets cytosolic misfolded proteins following heat stress. Nat Cell Biol 16, 1227–1237

89. Perez Berrocal, D. A., Witting, K. F., Ovaa, H., and Mulder, M. P. C. (2019) Hybrid Chains: A Collaboration of Ubiquitin and Ubiquitin-Like Modifiers Introducing Cross-Functionality to the Ubiquitin Code. Front Chem 7, 931

90. Anania, V. G., Pham, V. C., Huang, X., Masselot, A., Lill, J. R., and Kirkpatrick, D. S. (2014) Peptide level immunoaffinity enrichment enhances ubiquitination site identification on individual proteins. Mol Cell Proteomics 13, 145–156

91. Denison, C., Kirkpatrick, D. S., and Gygi, S. P. (2005) Proteomic insights into ubiquitin and ubiquitin-like proteins. Curr Opin Chem Biol 9, 69–75

92. Singh, R. K., Sundar, A., and Fushman, D. (2014) Nonenzymatic rubylation and ubiquitination of proteins for structural and functional studies. Angew Chem Int Ed Engl 53, 6120–6125

93. Verma, R., Peters, N. R., D’Onofrio, M., Tochtrop, G. P., Sakamoto, K. M., Varadan, R., Zhang, M., Coffino, P., Fushman, D., Deshaies, R. J., and King, R. W. (2004) Ubistatins inhibit proteasome-dependent degradation by binding the ubiquitin chain. Science 306, 117–120

94. Nakasone, M. A., Lewis, T. A., Walker, O., Thakur, A., Mansour, W., Castaneda, C. A., Goeckeler-Fried, J. L., Parlati, F., Chou, T. F., Hayat, O., Zhang, D., Camara, C. M., Bonn, S. M., Nowicka, U. K., Krueger, S., Glickman, M. H., Brodsky, J. L., Deshaies, R. J., and Fushman, D. (2017) Structural Basis for the Inhibitory Effects of Ubistatins in the Ubiquitin-Proteasome Pathway. Structure 25, 1839–1855 e1811

95. Dick, L. R., and Fleming, P. E. (2010) Building on bortezomib: second-generation proteasome inhibitors as anti-cancer therapy. Drug discovery today 15, 243–249

96. Jin, B., Wang, J., Liu, X., Fang, S., Jiang, B., Hofmann, K., Yin, J., and Zhao, B. (2018) Ubiquitin-Mimicking Peptides Transfer Differentiates by E1 and E2 Enzymes. Biomed Res Int 2018, 6062520

97. Thrower, J. S., Hoffman, L., Rechsteiner, M., and Pickart, C. M. (2000) Recognition of the polyubiquitin proteolytic signal. EMBO J 19, 94–102

98. Singh, S. K., Sahu, I., Mali, S. M., Hemantha, H. P., Kleifeld, O., Glickman, M. H., and Brik, A. (2016) Synthetic Uncleavable Ubiquitinated Proteins Dissect Proteasome Deubiquitination and Degradation, and Highlight Distinctive Fate of Tetraubiquitin. J Am Chem Soc 138, 16004–16015

99. Galan, J. M., and HaguenauerTsapis, R. (1997) Ubiquitin Lys63 is involved in ubiquitination of a yeast plasma membrane protein. Embo Journal 16, 5847–5854

100. Lauwers, E., Erpapazoglou, Z., Haguenauer-Tsapis, R., and Andre, B. (2010) The ubiquitin code of yeast permease trafficking. Trends Cell Biol 20, 196–204

101. Silva, G. M., Finley, D., and Vogel, C. (2015) K63 polyubiquitination is a new modulator of the oxidative stress response. Nat Struct Mol Biol 22, 116–123

102. Fang, N. N., Zhu, M., Rose, A., Wu, K. P., and Mayor, T. (2016) Deubiquitinase activity is required for the proteasomal degradation of misfolded cytosolic proteins upon heat-stress. Nat Commun 7, 12907

103. Isasa, M., Suner, C., Diaz, M., Puig-Sarries, P., Zuin, A., Bichman, A., Gygi, S. P., Rebollo, E., and Crosas, B. (2016) Cold Temperature Induces the Reprogramming of Proteolytic Pathways in Yeast. J Biol Chem 291, 1664–1675

104. Burroughs, A. M., Iyer, L. M., and Aravind, L. (2012) The natural history of ubiquitin and ubiquitin-related domains. Front Biosci (Landmark Ed) 17, 1433–1460

105. Song, L., and Luo, Z. Q. (2019) Post-translational regulation of ubiquitin signaling. J Cell Biol 218, 1776–1786

106. Gruhler, A., Olsen, J. V., Mohammed, S., Mortensen, P., Faergeman, N. J., Mann, M., and Jensen, O. N. (2005) Quantitative phosphoproteomics applied to the yeast pheromone signaling pathway. Mol Cell Proteomics 4, 310–327

107. Yu, Z., Kleifeld, O., Lande-Atir, A., Bsoul, M., Kleiman, M., Krutauz, D., Book, A., Vierstra, R. D., Hofmann, K., Reis, N., Glickman, M. H., and Pick, E. (2011) Dual function of Rpn5 in two PCI complexes, the 26S proteasome and COP9 signalosome. Mol Biol Cell 22, 911–920

108. Huang, D. T., and Schulman, B. A. (2005) Expression, purification, and characterization of the E1 for human NEDD8, the heterodimeric APPBP1-UBA3 complex. Methods Enzymol 398, 9–20

109. Matiuhin, Y., Kirkpatrick, D. S., Ziv, I., Kim, W., Dakshinamurthy, A., Kleifeld, O., Gygi, S. P., Reis, N., and Glickman, M. H. (2008) Extraproteasomal Rpn10 restricts access of the polyubiquitin-binding protein Dsk2 to proteasome. Mol Cell 32, 415–425

110. Kirkpatrick, D. S., Gerber, S. A., and Gygi, S. P. (2005) The absolute quantification strategy: a general procedure for the quantification of proteins and post-translational modifications. Methods 35, 265–273

111. Longworth, J., and Dittmar, G. (2019) Assessment of Ubiquitin Chain Topology by Targeted Mass Spectrometry. Methods Mol Biol 1977, 25–34

112. Mendes, M. L., Fougeras, M. R., and Dittmar, G. (2020) Analysis of ubiquitin signaling and chain topology cross-talk. Journal of proteomics 215, 103634

113. Kirkpatrick, D. S., Hathaway, N. A., Hanna, J., Elsasser, S., Rush, J., Finley, D., King, R. W., and Gygi, S. P. (2006) Quantitative analysis of in vitro ubiquitinated cyclin B1 reveals complex chain topology. Nat Cell Biol 8, 700–710

114. Mirzaei, H., Rogers, R. S., Grimes, B., Eng, J., Aderem, A., and Aebersold, R. (2010) Characterizing the connectivity of poly-ubiquitin chains by selected reaction monitoring mass spectrometry. Mol Biosyst 6, 2004–2014

115. Weber, A., Cohen, I., Popp, O., Dittmar, G., Reiss, Y., Sommer, T., Ravid, T., and Jarosch, E. (2016) Sequential Poly-ubiquitylation by Specialized Conjugating Enzymes Expands the Versatility of a Quality Control Ubiquitin Ligase. Mol Cell 63, 827–839

116. MacLean, B., Tomazela, D. M., Shulman, N., Chambers, M., Finney, G. L., Frewen, B., Kern, R., Tabb, D. L., Liebler, D. C., and MacCoss, M. J. (2010) Skyline: an open source document editor for creating and analyzing targeted proteomics experiments. Bioinformatics 26, 966–968

