## Supplemental Figures and Tables for "Rub1/NEDD8, a ubiquitin-like modifier, is also a ubiquitin modifier"

**Running title:** ***Rub1 modifies ubiquitin***

**This File contains:**

**Eight supplementary figures S1-S8** **P. 2-11**

**Three supplementary Tables S1-S3** **p. 12-13**

**Fig S1**

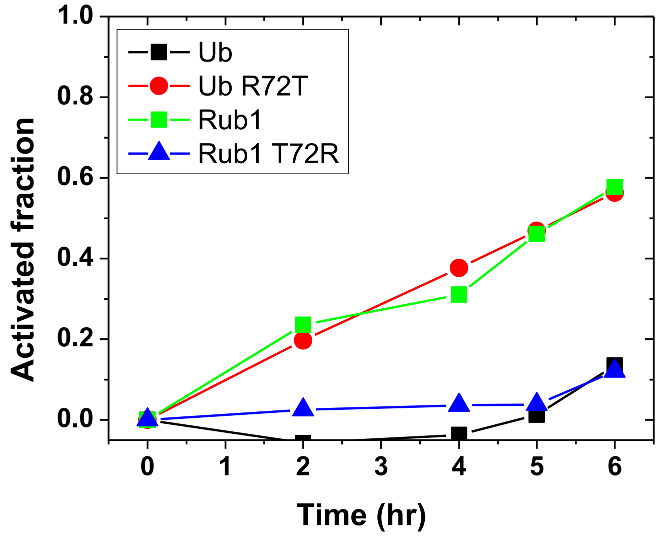


**Supplemental Figure S1 (For Figure 1):**

**Comparison of Ub and Rub1 activation by the NEDD8-activating enzyme, NAE.**

0.5 mM of recombinant Ub, Rub1, or their variants at position 72 were incubated with 0.5 µM of NAE at 30 $^{0}$ C in presence of 0.1 M MESNa to covalently label Ub/Rub1 thioesterified by E1. Fraction of Ub or Rub1 activated by UBA1 was assessed by quantifying residual inactivated protein as a function of the incubation time using an MS-based method (1).

*Method for Supplemental Figure S1*

*Recombinant Enzyme*: Human NAE (pGST-E1-NEDD8 from Brenda Schulman) was expressed as GST fusion construct in BL21(DE3) cells and purified using GST affinity columns as described (2)(3).

*Mass spectrometry assay of Ub / Rub1 activation by E1 enzymes***.** The amount of Ub or Rub1 (and variants) activated by the E1 enzyme was assessed using the MS-based method detailed in (1). Briefly, 0.5 mM of pure (unlabeled) Ub or Rub1 was incubated with 0.5 µM of respective E1 at 30 $^{0}$C along with 10 mM ATP, 10 mM $MgCl_2$, and 0.1 M MESNa in a 20 mM sodium-phosphate buffer at pH 8.0. All concentrations indicated here are the final concentrations of the components in a total of 100 µL reaction mixture. Equal amounts of reactions (20 µL) were aliquoted and stopped by adding 4 µL of 0.5 M EDTA. An internal standard of Ub or Rub1 was added to a final concentration of 0.5 mM. The aliquots were buffer exchanged with water (3X) using 3 kDa cut-off filters (Amicon, 0.5 mL volume). 10 µL of samples were mixed with 2 µL of 0.4% (v/v) TFA and injected (3X) into a mass spectrometer in ESI+ flight mode. $^{15}$N-enriched Ub was used as internal standard for measuring activation of Ub and all Ub variants by UAE or NAE. For Rub1 activation assays we used Rub1$^{T72R}$ for Rub1 activation by UAE or NAE, and Rub1 for Rub1$^{T72R}$ activation by UAE or NAE.

*Quantitation of Ub or Rub1 activation by E1***:** The activated fraction of Ub or Rub1 (or their variants) was obtained as follows. We monitored/measured the intensity ($I_t$) of the MS signal corresponding to the unactivated protein as a function of the incubation time. For accurate comparison of the $I_t$ values at different time points the MS signal intensities $I_t^*$ determined directly from the spectra were normalized/scaled using the intensity ($I^{st}$) of the internal standard signal in the same MS spectrum: $I_t = I_t^* \times I_0^{st} / I_t^{st}$ . The fraction of Ub or Rub1 thioesterified by E1 (further referred to as the "activated fraction") was quantified as follows:

activated fraction = $(I_0 - I_t)/I_0$,

where $I_0$ the initial/total MS signal of the protein (measured at time t=0 h), and $I_t$ is the normalized MS signal intensity corresponding to the remaining unactivated protein at a given time point *t*.

We note that we did not use/measure the intensity of the activated signal directly because (a) it has a different charge from the unactivated protein and therefore could have an altered time of flight in mass spectrometry, and (b) it is not always discernible. The plot shows the activated fraction (which is =1-($I_t/I_0$) ) on a scale from 0 to 1; if multiplied by 100 it would give the percentage of protein activated.

**Fig S2**

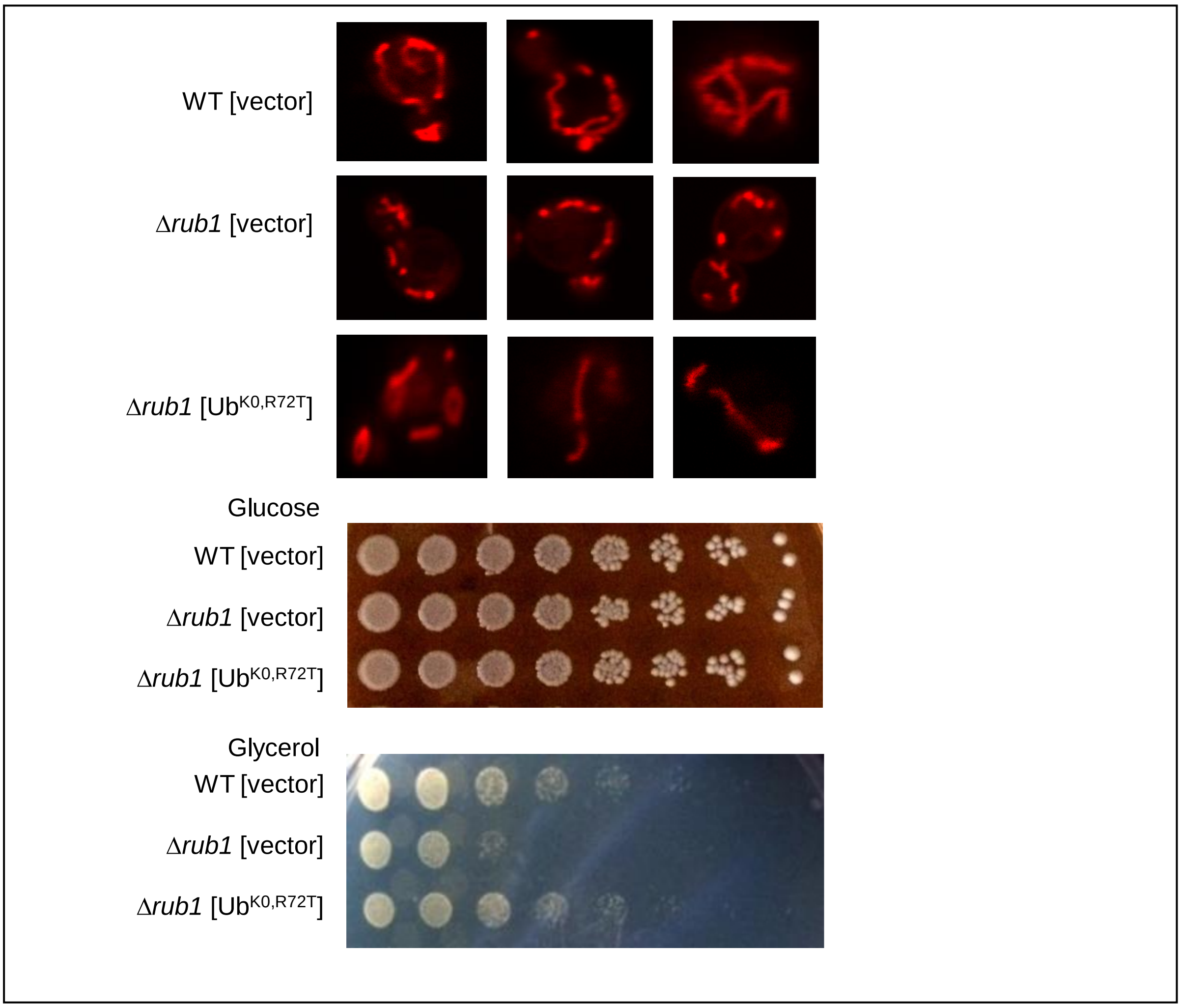


**Supplementary Figure S2 (for Figure 2):**

**Lysineless ubiquitin reverses mitochondrial phenotypes associated with loss of *RUB1*.**

Top. Wild-type *S. cerevisiae,* Δ*rub1*, or Δ*rub1* expressing Ub$^{K0,R72T}$ were treated with MitoTracker and mitochondria monitored by confocal fluorescence microscopy (LSM700).

Bottom. Cultures of Wild-type *S. cerevisiae,* Δ*rub1*, or Δ*rub1* expressing Ub$^{K0,R72T}$ were serially diluted onto selective glucose-containing media (top) or Glycerol (bottom) and grown at 30C for 3 days.

Method: *Fluorescent microscopy for mitochondria*. Cells in exponential growth phase were collected by centrifugation, and 1 µM MitoTracker Red was added for 30 minutes. Cells were washed twice and the mitochondria in cells were monitored using LSM700 confocal microscope.

**Fig S3**

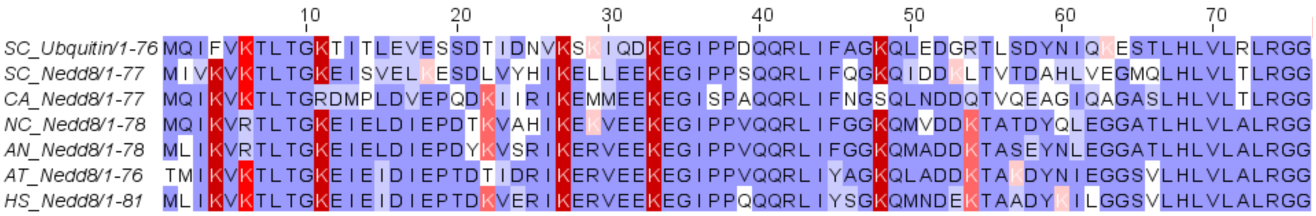


**Supplementary Figure S3 (for Figure S3):**

**Multiple sequence alignment for Rub1/Nedd8 from various species**.

Alignment of *Candida albicans* Rub1 protein sequence with other Rub1/Nedd8 and ubiquitin. Lysines are highlighted in red (intensity reflecting conservation). Residues diverging from the consensus are highlighted in white. Note that the Rub1 sequence in *C. albicans* lacks Lysines at positions 11, 48 and 63, which generate the three most abundant ubiquitin linkages according to quantification in various cell types. All Rub1/NEDD8 sequences differ from the invariant arginine of ubiquitin at position 74.

SC- *Saccharomyces cerevisiae*, CA- *Candida albicans*, NC- *Neurospora crassa*, AN- *Aspergillus nidulans*, AT- *Arabidopsis thaliana*, HS- *Homo sapiens*.

Method: Ubiquitin and Rub1/Nedd8 sequences of different organisms were collected from NCBI protein database (https://www.ncbi.nlm.nih.gov/protein). Sequences were aligned using CLUSTALW multiple sequence alighnment tool and was edited and visualized in JALVIEW.

**Fig S4**

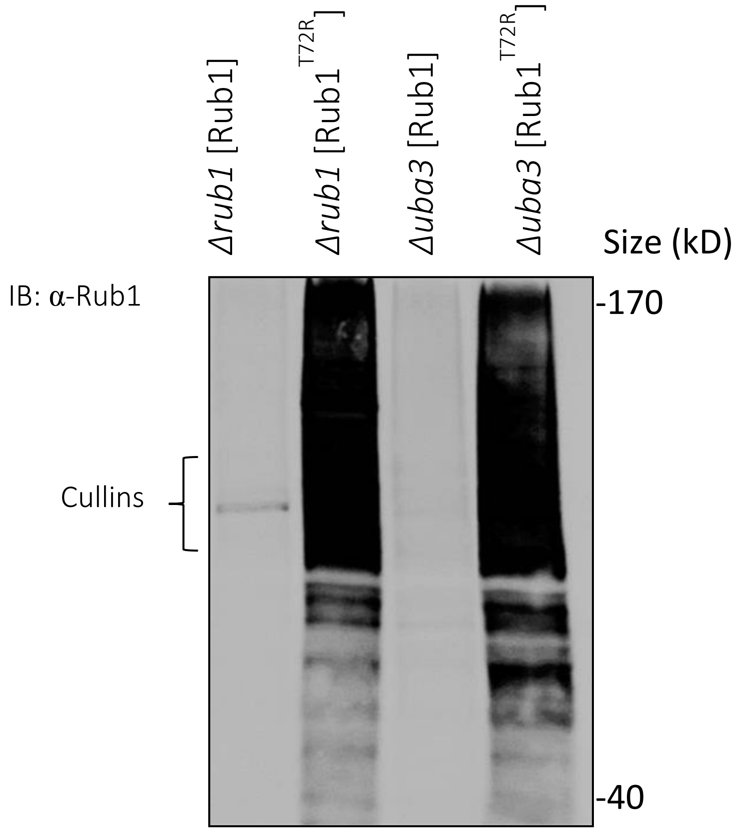


**Supplementary Figure S4 (For Figure 4):**

**Rubylation landscape of Rub1$^{T72R}$ independent of NAE**. Δ*rub1* or Δ*uba3* complemented by Rub1 or Rub1$^{T72R}$ as noted were grown to mid log phase, lysed, and whole cell extract resolved by 8% SDS PAGE and immunoblotted for Rub1. Prominent bands at ~100 KDa in WT (left) reflect Rub1-modified cullins, which are undetectable in absence of NAE (third lane). Myriad targets are detected i=with "Ubiqutinized" Rub1 (Rub1$^{T72R}$) independent of functional NAE (right lane).

**Fig S5**

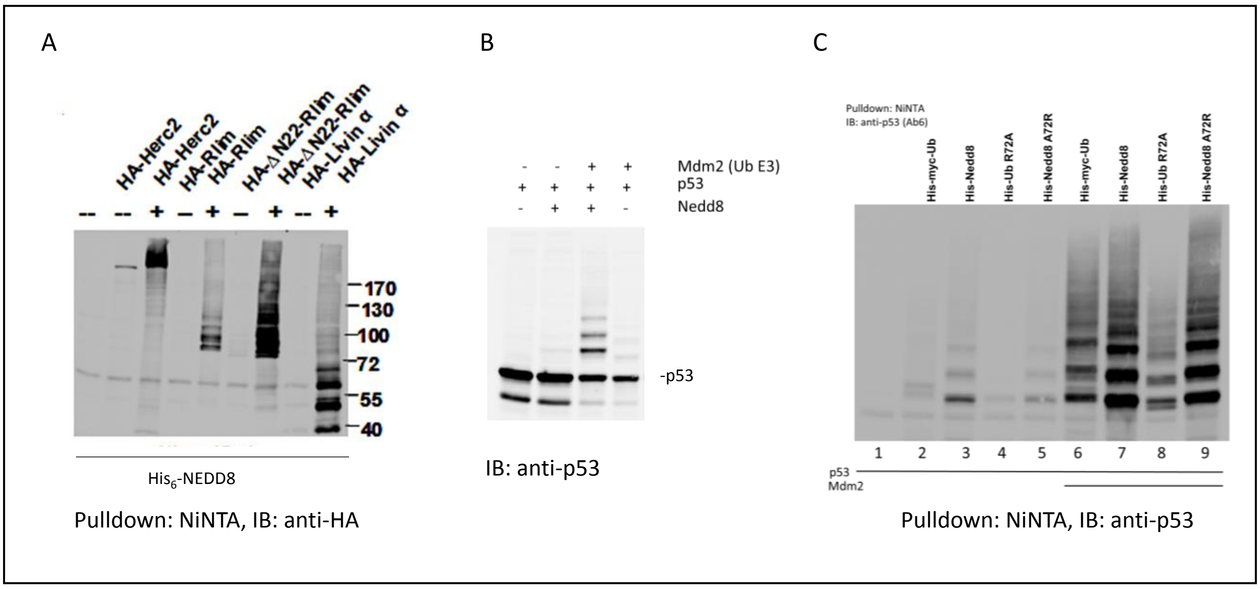


**Supplementary Figure S5 (For Figure 4):**

**Overexpressed NEDD8 in mammalian cells leads to NEDDylation of substrates for ubiquitination.**

A. 1 µg of each plasmid, as indicated, was transfected into H1299 cells. 20-24 hours posttransfection, cells were lysed in denaturing guanidinium hydrochloride buffer. His-Nedd8 conjugated proteins were purified by Ni-beads followed by SDS-PAGE and western blot performed with indicated antibodies.

B. H1299 cells were transfected with 200 ng of each plasmid mixed with 200 ng of β-galactosidase plasmid, except for NEDD8 plasmid that was transfected at 1 µg. Cells were harvested 20-24 hours post-transfection and lysed in TNN buffer. Transfection efficiency was normalized by β-galactosidase assay followed by SDS-PAGE and Western blot.

C. The plasmids His-Ub or its mutants and His-Nedd8 or its mutants, 1 µg each were transfected into H1299 cells. After 20-24 hours of transfection, cells were lysed in denaturing guanidinium hydrochloride buffer. p53 conjugated with His tagged proteins were purified by Ni-beads followed by SDS-PAGE and western blot performed with a HA antibody. The difference in the positions for wt-Ub modified p53 and the Ub mutant modified p53 on the blot was due to the fact that wt-Ub had a His-myc tag whereas its mutant had only the His tag.

**Fig S6**

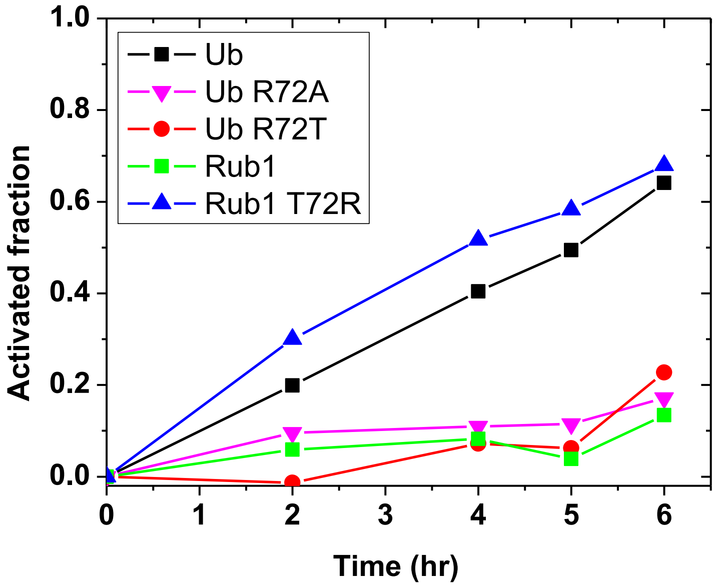


**Supplementary Figure S6 (For Figure 4):**

**Comparison of Ub and Rub1 activation by the E1 Ub-activating enzyme, UAE.**

0.5 mM of recombinant Ub, Rub1, or their variants at position 72 were incubated with 0.5 µM of NAE (UBA3-NAE1 E1 enzyme) at 30 $^0$ C in presence of 0.1 M MESNa to covalently label Ub/Rub1 thioesterified by E1. Fraction of Ub or Rub1 activated by UBA1 was assessed by quantifying residual inactivated protein as a function of the incubation time using an MS-based method (1).

Method – as in supplementary Figure S1.

**Fig S7**

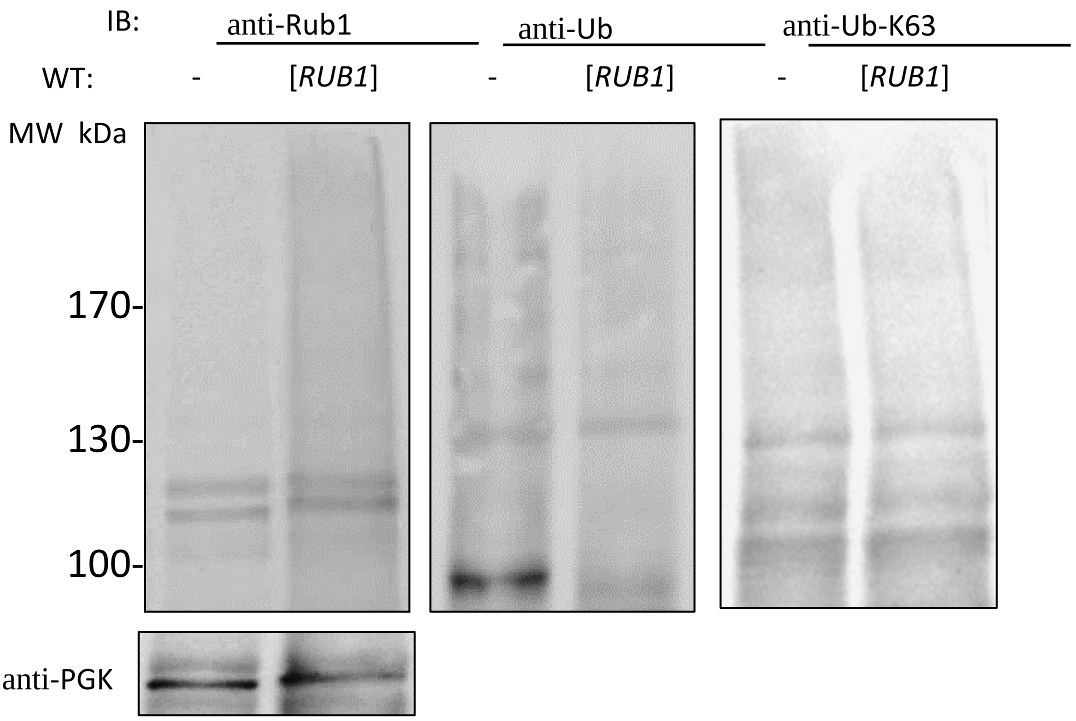


**Supplementary Figure S7 (For Figure 4**):

**Introducing Rub1 in the Ubiquitin landscape.**

Whole cell extracts from log phase WT, or WT overexpressing *RUB1* resolved by 8% SDS PAGE and immuno-blotted for Rub1, Ub or K63-Ub linkages, respectively.

**Fig S8**

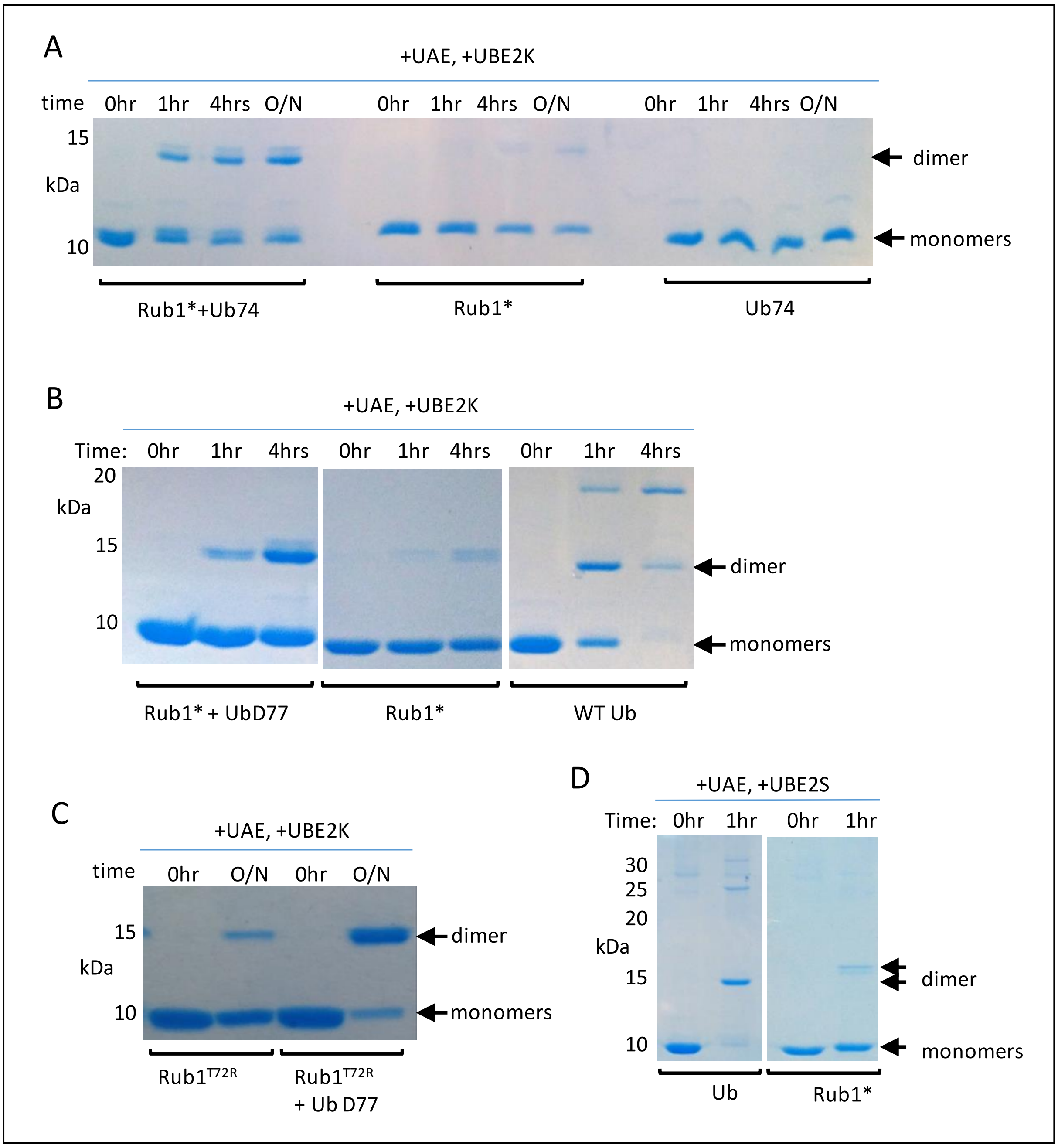


**Supplementary Figure S8 (For Figure 5)**:

**Ubiquitin E2 conjugating enzymes differentiate between Ub and Rub1.**

A. Equimolar Rub1* ("ubiqutinized" at position 72 to render it a substrate for UBA1) and Ub74 (truncated at position 74 to render it inert to activation by UBA1) were incubated up to 12 hours together or separately with the Ub E1 enzyme, UBA1, and the Ub E2 enzyme, UBE2K. Migration of the monomeric substrates and product dimer are marked by arrows on right. Note: we label Rub1$^{T72R}$ in this panel as Rub1* since sequencing identified an accompanying mutation H60N,T72R. B. Similar reaction as in A, but with an extended ubiquitin specie, UbD77 that also renders it inert to UBA1 activation (left) or with wild-type ubiquitin the preferred substrate for UBA1 that together with UBE2K generates predominantly K48-linked dimers, trimers and higher order polymeric ubiquitin chains (right). C. Similar reaction as in B, incubated for 12 hours. D. ("ubiqutinized" Rub1$^{T72R}$ or Ub were incubated for 1 hour with the Ub E1 enzyme, UBA1, and the Ub E2 enzyme, UBE2S. Migration of the monomeric substrates and product dimer are marked by arrows on right. UBE2S generates predominantly K11-linked dimers, trimers and higher order polymeric ubiquitin chains (left).

Method for supplementary figure S8 - *Enzymatic assembly of Rub1-Ub conjugates*. WT or mutated Rub1 and Ub (10 mg each) were incubated with 20-µM ubiquitin-conjugating E2 enzyme (UBE2K, aka E2-25 K), 500-nM ubiquitin-activating E1 enzyme (UBE1, aka UBA1), 10 mM creatine phosphate, 5 mM $MgCl_2$, 5 mM ATP, and creatine phosphokinase in 50 mM Tris-HCl buffer (pH 8) at room temperature for approximately 16 hours. The solution was centrifuged at 13 000 rpm in order to remove the precipitated enzymes (E1 and E2) and further fractionated using FPLC.

**Supplementary Table S1: Strains used in this study**

| Strain | Phenotype | Background | Genotype | Source |
|---|---|---|---|---|
| | | *S.cerevisiae* | | |
| MY58 | WT | BY4741 | MATa; his3ko1; leu2ko0; met15ko0; ura3ko0 | EUROSCARF |
| MY299 | ΔRUB1 | BY4741 | MATa; his3ko1; leu2ko0; met15ko0; ura3ko0; YDR139c::kanMX4 | UROSCARF |
| MY101 | ΔCSN5 | BY4742 | MATα; his3ko1; leu2ko0; lys2ko0; ura3ko0; DYDL216c:hisMX6; CSN5::kanMX4 | EUROSCARF |
| MY1297 | ΔRUB1ΔCSN5 | BY4742 | MATα; his3ko1; leu2ko0; lys2ko0; ura3ko0; DYDL216c:hisMX6; YDR139c::kanMX4 | EUROSCARF |
| MY302 | ΔUBA3 | BY4742 | MATα; his3ko1; leu2ko0; lys2ko0; ura3ko0; DYDL216c:hisMX6; UBA3::kanMX4 | EUROSCARF |
| MY878 | Cdc53 K760R | BY4741 | MATa; his3ko1; leu2ko0; lys2ko0; met15ko0; ura3ko0; YDL132w::kanMX4; Leu-cdc53 K760R | EUROSCARF |
| | | *C. albicans* | | |
| KC208 | *rub1-/-* | | ura3-/ura3-, rub1-/rub1- | (4) |

**Supplementary Table S2: Plasmids used in this study**

| Plasmid | description | Source |
|---|---|---|
| m201 | Backbone Yeplac plasmid with ADH promotor | |
| m1564 | RGS-His$_8$-WT Rub1 under ADH1 promoter | This study |
| m1532 | RGS-His$_8$-Rub1 T72R | This study |
| m886 | RGS-His$_8$-WT Ub under ADH1 promoter | This study |
| m965 | RGS-His$_8$-K0 Ub under ADH1 promoter | This study |
| m1540 | RGS-His$_8$-Ub R72T under ADH1 promoter | This study |
| m1541 | RGS-His$_8$-K0 Ub R72T under ADH1 promoter | This study |
| m699 | Cdc53 K760R under native promoter | This study |
| | pTXB1-Rub1 (Rub1) | (5) |
| | pTXB1-Rub1 T72R (Rub1$^{T72R}$) | (5) |
| | pTXB1-Rub1 E63K and T72R (Rub1$^{E63K,T72R}$) | (5) |
| | pTXB1-Rub1 H60N and T72R (Rub1$^{H60N,T72R}$) | (5) |

**Supplementary Table S3: SRM conditions for the unlabeled ubiquitin peptides**

| Chain type | Name of the pept | Sequence | Q1 mass | Q3 mass | DP(V) | CE(V) |
|---|---|---|---|---|---|---|
| K48 | | LIFAGK(GG)QLEDGR | 487.6 | 617.8<br>347.2 | 130 | 19<br>33 |
| K63 | | TLSDYNIQK(GG)ESTLHLVLR | 748.7 | 1015.5<br>1067.6 | 130 | 32<br>46 |
| ubiquitin | Ubiquitin total | EGIPPDQQR | 520.3 | 740.4<br>643.3 | 130 | 29<br>30 |
| | 14 ubiquitin_#1 | ESTLHLVLR | 356.5 | 431.2<br>426.3 | 130 | 17<br>17 |
| | 15 ubiquitin_#2 | TLSDYNIQK | 541.3 | 867.4<br>665.4 | 130 | 29<br>31 |

### References

1. Singh, R. K., Kazansky, Y., Wathieu, D., and Fushman, D. (2017) Hydrophobic Patch of Ubiquitin is Important for its Optimal Activation by Ubiquitin Activating Enzyme E1. *Anal Chem* **89**, 7852-7860
2. Singh, R. K., Zerath, S., Kleifeld, O., Scheffner, M., Glickman, M. H., and Fushman, D. (2012) Recognition and cleavage of related to ubiquitin 1 (Rub1) and Rub1-ubiquitin chains by components of the ubiquitin-proteasome system. *Mol Cell Proteomics* **11**, 1595-1611
3. Huang, D. T., and Schulman, B. A. (2005) Expression, purification, and characterization of the E1 for human NEDD8, the heterodimeric APPBP1-UBA3 complex. *Methods Enzymol* **398**, 9-20
4. Sela, N., Atir-Lande, A., and Kornitzer, D. (2012) Neddylation and CAND1 independently stimulate SCF ubiquitin ligase activity in Candida albicans. *Eukaryot Cell* **11**, 42-52
5. Gomes, F., Lemma, B., Abeykoon, D., Chen, D., Wang, Y., Fushman, D., and Fenselau, C. (2019) Top-down analysis of novel synthetic branched proteins. *J Mass Spectrom* **54**, 19-25
